# Single Cell Resolution Spatial Mapping of Human Hematopoiesis Reveals Aging-Associated Topographic Remodeling

**DOI:** 10.1101/2023.04.28.538715

**Authors:** Aleksandr Sarachakov, Arina Varlamova, Viktor Svekolkin, Ilia Galkin, Itzel Valencia, Caitlin Unkenholz, Tania Pannellini, Aida Akaeva, Sofia Smirnova, Pavel Ovcharov, Margarita Polyakova, Dmitrii Tabakov, Ekaterina Postovalova, Isha Sethi, Nara Shin, Alexander Bagaev, Tomer Itkin, Genevieve Crane, Michael Kluk, Julia Geyer, Giorgio Inghirami, Sanjay Patel

**Author notes:** Correspondence: Sanjay S. Patel, MD.

## Abstract

The spatial anatomy of hematopoiesis in bone marrow has been extensively studied in mice and other preclinical models, but technical challenges have precluded a commensurate exploration in humans. Institutional pathology archives contain thousands of paraffinized bone marrow core biopsy tissue specimens, providing a rich resource for studying the intact human bone marrow topography in a variety of physiologic states. Thus, we developed an end-to-end pipeline involving multiparameter whole tissue staining, in situ imaging at single-cell resolution, and artificial intelligence (AI)-based digital image analysis, and then applied it to a cohort of disease-free samples to survey alterations in the hematopoietic topography associated with aging. Our data indicate heterogeneity in marrow adipose tissue (MAT) content within each age group, and an inverse correlation between MAT content and proportions of early myeloid and erythroid precursors, irrespective of age. We identify consistent endosteal and perivascular positioning of hematopoietic stem and progenitor cells (HSPCs) with medullary localization of more differentiated elements and, importantly, uncover new evidence of aging-associated changes in cellular and vascular morphologies, microarchitectural alterations suggestive of inflammaging, and diminution of a potentially active megakaryocytic niche. Overall, our findings suggest that there is topographic remodeling of human hematopoiesis associated with aging. More generally, we demonstrate the potential to deeply unravel the spatial biology of normal and pathologic human bone marrow states using intact archival tissue specimens.

## INTRODUCTION

Hematopoiesis is the tightly-coordinated process within mammalian bone marrow that results in the continuous repopulation of the blood’s cellular constituents throughout the lifetime of an organism.^1^ Numerous studies over many decades have consistently demonstrated the importance of spatially distinct haematopoietic “niches”, regulatory units comprised of hematopoietic stem and progenitor cells (HSPCs), non-hematopoietic cellular elements, and critical structural co-constituents,^2,3^ which govern HSPC activity and thereby maintain haematopoietic stability. Our current knowledge of the *intact* marrow topography has largely been informed by sophisticated imaging studies performed in recent decades, primarily using mice and other preclinical models.^2,4– 16^ Based on these studies, at least two anatomically distinct niches have been identified and rigorously studied: 1) the endosteal niche,^4,5,9,17–23^ in close proximity to the epiphyseal and diaphyseal/trabecular bone surfaces, and 2) the central or vascular niche, which contains the majority of the sinusoids and arterioles located throughout the marrow.^6,8–10,12,24–34^ In addition, some studies have provided evidence that other marrow constituents such as megakaryocytes,^14,35–38^ macrophages,^39–41^ and adipocytes^42^ may also influence hematopoietic stem cell (HSC) biology.

The mouse has proven to be an indispensable model system for studying hematopoiesis, but there remains a relative paucity of single cell, and multiparameter in situ imaging data, generated using samples from humans or non-human primates. Whether or not murine models faithfully recapitulate the human state remains largely unknown.^43^ Moreover, most bone marrow imaging studies in mice have used femoral long bones rich in red marrow, while adult human femurs are rich in yellow marrow with limited hematopoiesis. These and other differences may explain why preclinical findings have infrequently translated into effective clinical interventions;^44^ however, technical and ethical challenges have historically prohibited modeling hematopoiesis in humans. It is therefore not surprising that very few studies to date have attempted to examine bone marrow topography using human tissue, and have been restricted by small cohort sizes, a limited number of analyzed parameters, and primarily visual or only semi-quantitative analytical techniques.^45–52^ Institutional pathology archives can contain many thousands of paraffinized bone marrow core biopsy specimens that remain underutilized following initial diagnostic evaluations and therefore represent a rich resource with which to study the intact human bone marrow topography in a variety of states and additionally model disease longitudinally in individual patients. Recent advances in tissue imaging^53–58^ and computational bioinformatics have enabled a more robust assessment of such samples, creating an opportunity to study the marrow topography in humans and translate this knowledge into promising and efficacious therapies for patients.

Here, we describe an end-to-end pipeline to study archival bone marrow samples, involving multiparameter whole tissue staining, whole slide in situ imaging at single-cell resolution, and artificial intelligence (AI)-based digital image analysis. We further demonstrate the application of this pipeline to survey changes in the topography of human hematopoiesis associated with aging.

## RESULTS

### Multiparameter whole tissue staining, in situ imaging, and AI-driven image analysis of human bone marrow core biopsy tissues at single-cell resolution

Individuals who underwent a posterior iliac crest bone marrow biopsy evaluation, either as part of a staging procedure, or to evaluate for a mild cytopenia of unknown etiology, were considered for inclusion in the study. Young (<20 years, n=11), middle-aged (20-60 years, n=5), and older (>60 years, n=13) individuals without evidence of a bone marrow pathology on comprehensive hematopathology evaluation, including cytogenetic analysis, were selected. The cohort included 16 men and 13 women. Accompanying laboratory data were collected, including complete blood count (CBC) parameters such as hemoglobin and mean corpuscular volume (MCV) values, absolute neutrophil (ANC) and platelet counts, mean platelet volumes, and marrow aspirate smear differential counts, and orthogonal flow cytometry data for a subset of cases (**Supplemental Data Table 1**). Hematoxylin and eosin (H&E)-stained preparations of the bone marrow core biopsy tissue specimens were first re-evaluated by light microscopy to determine adequacy and quality of remaining paraffin-embedded tissue. Serial consecutive sections from qualified samples were then prepared by multiplex immunofluorescence (MxIF) staining in an automated tissue stainer, followed by whole slide imaging, spectral unmixing, and feature extraction (**Figure 1A**). The entire bone marrow tissue area in each sample was analyzed, including individual cell detection, cell segmentation and phenotyping, in conjunction with machine learning-based detection of bone trabeculae, CD34+ endothelial cell-lined vasculature, and marrow adipose tissue (**Figure 1B, Supplemental Data Fig. 1**). A total of 1,510,295 nucleated cells were analyzed across the entire cohort. A convolutional neural network (CNN) approach was employed to classify cells based on morphologic features, mean fluorescence intensities (MFIs) of detected antigens, and antigen expression patterns (see Methods). A total of 11 different cell types were identified: hematopoietic stem and progenitor cells, myeloblasts, promyelocytes, proerythroblasts, maturing myeloid cells (MMCs), erythroid normoblasts, megakaryocytes, likely B-cell precursors (LBPs), likely plasma cells, mast cells, and non-hematopoietic elements (**Figure 1C**); CD34+ endothelial cells were also detected. Nucleated cells lacking expression of any of the interrogated antigens were classified as non-hematopoietic elements (NHEs). A generated vessel mask for each image (see Methods) was used to distinguish CD34+ HSPCs and CD34+ endothelial cells, by integrating MFI values and morphologic features. Percentages of various cell types quantified from MxIF images were correlated with orthogonal data generated by manual aspirate smear differential counting and/or multiparameter flow cytometric (MFC) immunophenotyping performed as part of the original diagnostic evaluation. We were able to identify a positive correlation between the myeloid to erythroid ratio, as determined by aspirate smear differential count, and the ratio of MMCs to erythroid normoblasts generated by MxIF analysis (Spearman R = 0.45, p=0.025) [**Supplemental Data Fig. 2**]; however, we did not observe significant correlations between quantification of individual cell types (e.g., myeloblasts, promyelocytes) between MxIF and orthogonal methods (p>0.05 for all tested correlations). These findings underscore the significance of assessing intact and unperturbed bone marrow tissue; aspirated marrow material used for morphologic evaluation, and other single-cell analyses, is composed of disaggregated cells that may not accurately reflect the true marrow contents and additionally cannot provide meaningful information about spatial cell-cell interactions or the entire marrow microenvironment.

**Figure 1.**
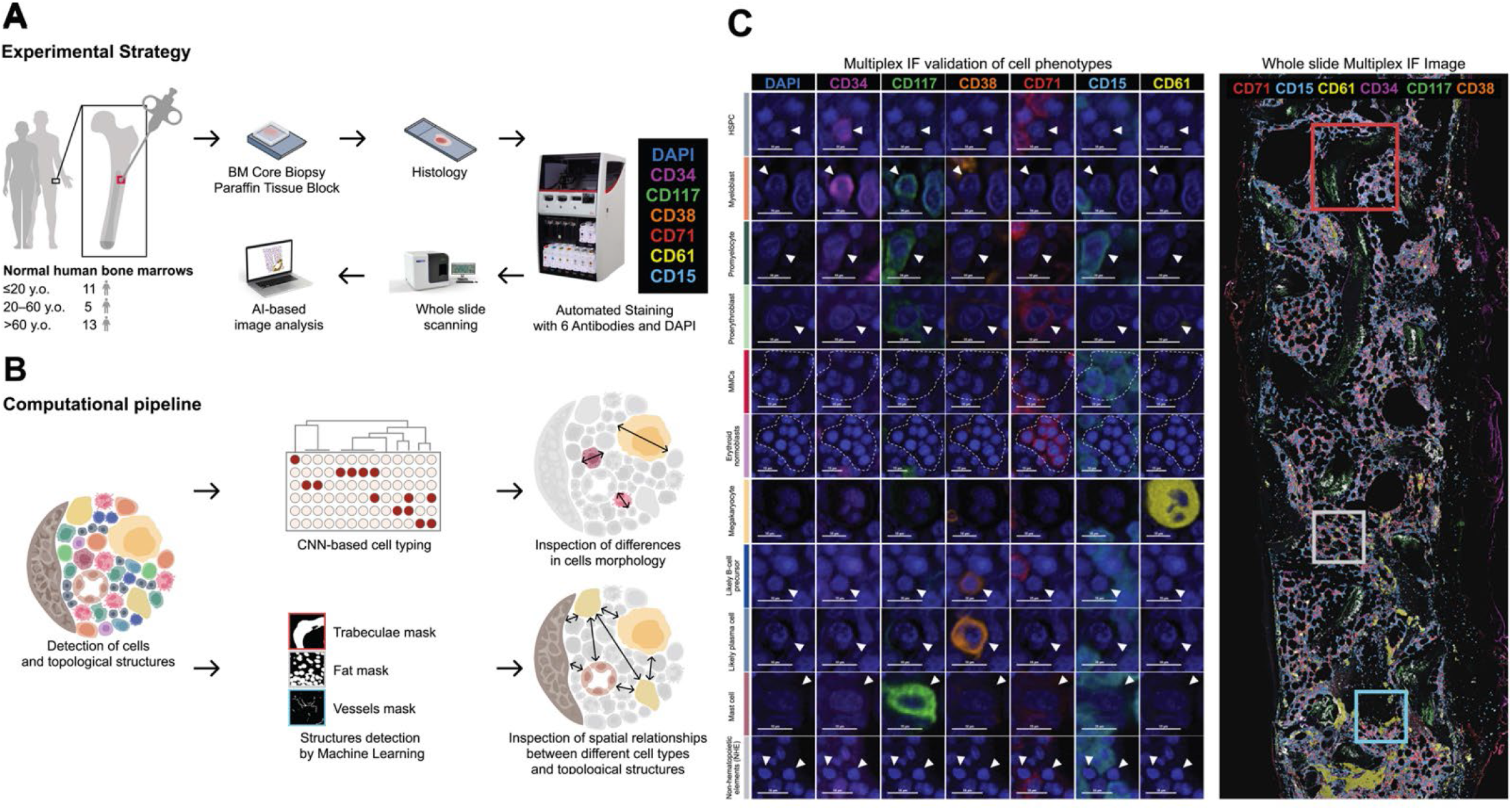
MxIF staining, whole tissue imaging, and AI-driven image analysis of human bone marrow core biopsy tissues at single cell resolution. **A**. Experimental strategy is shown. Archival normal bone marrow (BM) specimens from 29 individuals were collected. Paraffin-embedded BM core biopsy tissues were sectioned on to positively charged glass microscope slides. Automated iterative multiplex immunofluorescence staining was performed in a Bond RX autostainer (Leica Biosystems, Buffalo Grove, IL) with the following panel of antibodies plus DAPI: CD34, CD117, CD38, CD71, CD61, CD15. Whole slide images (WSIs) were captured with the Vectra Polaris Automated Quantitative Pathology Imaging System (Akoya Biosciences, Marlborough, MA). WSI tiles were spectrally unmixed in InForm (Akoya) and subsequently stitched in HALO (Indica Labs, Albuquerque, NM). Spectrally-unmixed WSIs were parsed through BostonGene-developed artificial intelligence (AI)/machine learning (ML)-based custom Python pipelines. **B**. Computational pipeline involving convolutional neural network (CNN)-based cell type identification (n=11) using marker mean fluorescence intensities (top); in parallel, machine learning-based masks for bone trabeculae, fat, and CD34+ vasculature were generated (bottom). Morphologic features of identified cell types (n=11) were assessed. Distances between different cell types and to various structures of interest were determined. **C**. Validation of detected cell types based on MxIF staining (left); cell types specified vertically and antibody channels listed horizontally. A total of 11 different cell types were identified: hematopoietic stem and progenitor cells (HSPCs) [median=1041/case, total=30463 cells, density=130 cells/mm^2^], myeloblasts (median=190/case, total=9102/case, density=39 cells/mm^2^), promyelocytes (median=246/case, total=10928/case, density=50 cells/mm^2^), proerythroblasts (median =199/case, total=10074/case, density=40 cells/mm^2^), maturing granulocytes and monocytes (maturing myeloid cells [MMCs], median=16500.0/case, total=597387 cells, density=2500 cells/mm^2^), erythroid normoblasts (median=11248/case, total=339276/case, density=1450 cells/mm^2^), megakaryocytes (median=372/case, total=12739 cells, density=60 cells/mm^2^), likely B-cell precursors (LBPs) [median=1979/case, total=71396/case, density=290 cells/mm^2^], and likely plasma cells (median=86.0/case, total=3520/case, density=10 cells/mm^2^). Nucleated cells lacking expression of any of the interrogated antigens were classified as non-hematopoietic elements (NHEs) [median=12682/case, total=392506/case, density=1740 cells/mm^2^]. Mast cells and CD34+ endothelial cells were also detected. MxIF WSI of a representative normal bone marrow biopsy is shown (right).

### Specific cell type proportions fluctuate with aging and marrow adiposity

Cell type proportions visually appeared consistent between age groups (**Figure 2A**). On inspection of individual subpopulations, we observed a significant reduction in the percentages of myeloblasts (0.8±0.5 vs. 0.4±0.25, p=0.01) and proerythroblasts (0.01±0.0035 vs. 0.0037±0.0027, p=0.00032) between the youngest and oldest individuals (**Figure 2B**). We also observed diminution of likely B-cell precursors (0.05±0.0198 vs. 0.039±0.0239, p=0.046) and mast cells (0.0259±0.0102 vs. 0.0134±0.0066, p=0.0054). Conversely, marrows from the oldest individuals had greater percentages of NHEs [0.2201±0.0493 vs. 0.3171±0.0876, p=0.00107] and endothelial cells (0.0013±0.0018 vs. 0.0029±0.0022, p=0.01603) [**Supplemental Data Fig. 3**]. We also identified significant negative correlations for percentages of the following cell types increasing age: myeloblasts, proerythroblast, maturing myeloid cells (MMCs), and mast cells; conversely, we found a significant positive correlation in the percentage of NHEs [**Figure 2B, Supplemental Data Fig. 3**]. We did not observe significant differences between age groups in the overall proportions of the other cell types evaluated, including HSPCs, promyelocytes, and megakaryocytes (p>0.05 for all). However, when we normalized for the total cell density (per mm^2^) in each case, we found that the normalized value for HSPCs was significantly greater in individuals 20-60 years compared to those ≤20 years (p<0.001) and >60 years (p<0.01) [**Figure 2B**]. This additional analysis suggests that HSPC content increases in the 20-60 year old group compared to younger individuals, but then begins to diminish in individuals >60 years of age.

**Figure 2.**
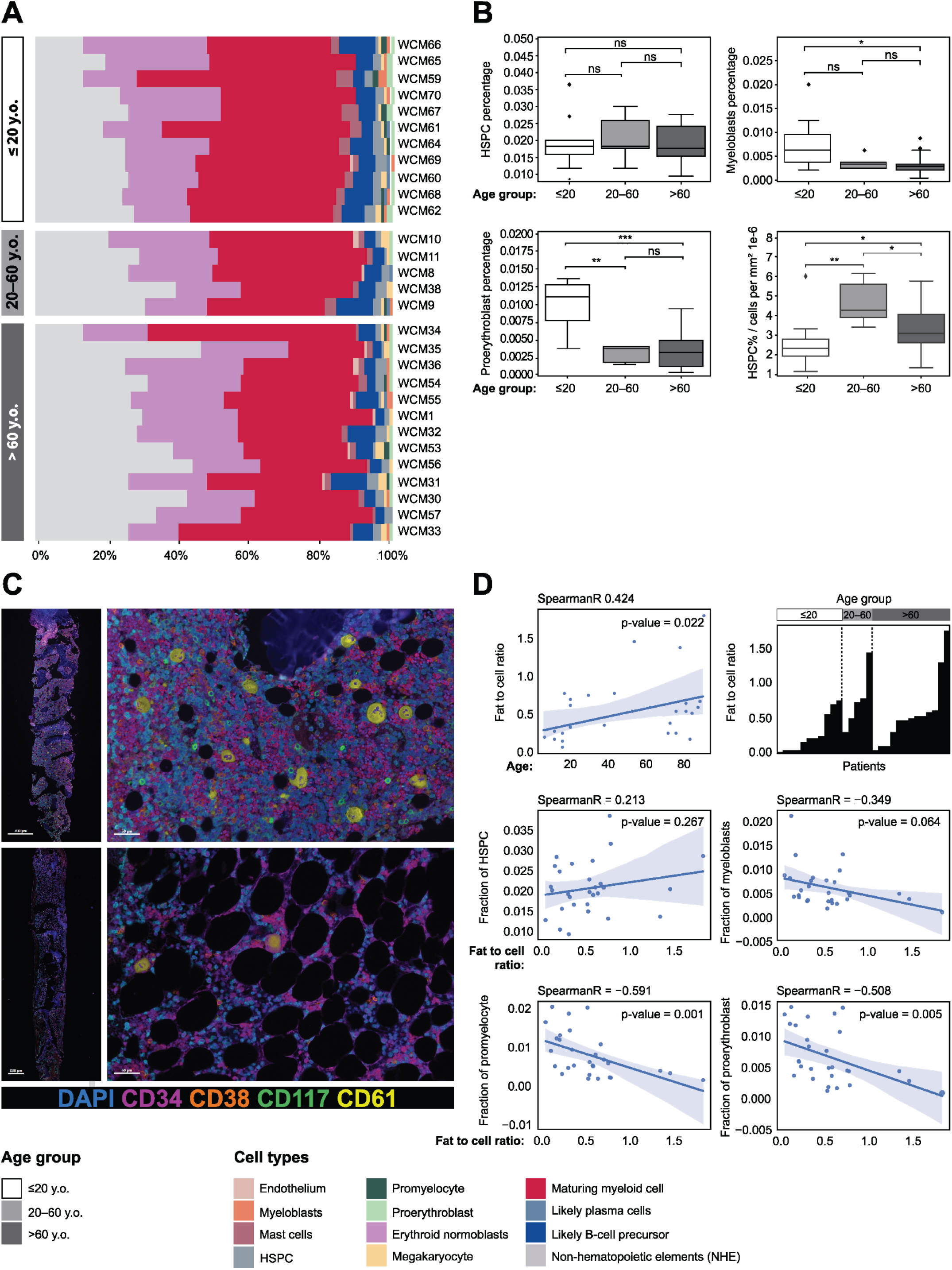
Alterations in cell type proportions associated with aging or adiposity. **A**. Relative proportions of hematopoietic and non-hematopoietic cell types depicted in each sample and grouped by age. **B**. Proportion of HSPCs does not differ with increasing age (p>0.05); however, an increase in proportion of HSPC is observed upon normalization to cell density/mm^2^ (2.57±1.3 vs 4.66±1.15, p=0.0055 for ≤20 years vs 20-60 years and 4.66±1.15 vs 3.34±1.19, p=0.0224 for 20-60 years vs >60 years; and 2.57±1.3 vs 3.34±1.19, p=0.0334 for ≤20 years vs >60 years). Pairwise comparisons between age groups for specific cell types shows diminished proportions of myeloblasts (0.0079±0.0053 vs. 0.0038±0.0026, p=0.01) and proerythroblasts (0.0104±0.0037 vs. 0.0037±0.0028, p=0.0003), between ≤20 years and >60 years BMs. Conversely, proportions of NHEs increased with aging (≤20 years and >60 years, 0.22±0.052 vs 0.317±0.09, p=0.01) as well as MMCs (≤20 years and >60 years, 0.42±0.07 vs 0.35±0.095, p=0.005). **C**. Representative MxIF examples of ≤20 years (WCM66, top) and >60 years (WCM57, bottom) BM specimens (right panel) show greater cellular density (top) and higher fat content (bottom), respectively. **D**. Correlation between fat:cell ratio and increasing age reveals a positive association (Spearman R = 0.424, p=0.022). Fat:cell ratio as quantified by AI-based image analysis shows heterogeneity within age groups with a trend toward increasing fat content in marrows from older individuals (top). Separation of the entire cohort at the median for fat:cell ratio reveals no significant difference in proportion of HSPCs (p>0.05). Increasing marrow adiposity correlates with diminishing proportions of myeloblasts (Spearman R = -0.349, p=0.064) and significant diminution of promyelocytes (Spearman R = -0.591, p=0.001) and proerythroblasts (Spearman R = -0.508, p=0.005), irrespective of age.

We observed an increasing fat-to-cell area ratio with older age (Spearman R = 0.424, p=0.022), although each patient age group included samples with a range of fat:cell proportions, indicating heterogeneity in cellular density irrespective of age (**Figure 2C-D**). Indeed, by groupwise analysis, a statistically significant difference in fat-to-cell area ratio was only observed between individuals ≤20 years and those between 20-60 years (0.21 vs. 0.56, p=0.038), but not between the extremes of age (p>0.05. Given the heterogeneity in fat-to-cell area ratio within age groups, we next asked whether higher fat density was associated with differences in cell type frequencies irrespective of patient age [**Supplemental Data Fig. 4**]. We found that the percentage of HSPCs did not differ significantly relative to marrow adiposity; however, marrows with higher fat-to-cell area ratios were associated with lower percentages of myeloblasts (Spearman R = -0.349, p=0.064) and significant diminution of promyelocytes (Spearman R = -0.591, p=0.001) and proerythroblasts (Spearman R = -0.508, p=0.005), suggesting that higher fat density may constrain initial erythroid and granulocytic output from HSPCs **[Figure 2D]**. Similar findings have been previously reported in mice.^59^ We did not identify differences in proportions of more mature elements such as MMCs or erythroid normoblasts (p>0.05 for both). Higher fat density marrows were associated with higher erythrocyte MCV values (81.9±8.2 vs. 88.5±8.0, p = 0.0137), although we did not observe significant differences in other CBC parameters (e.g. hemoglobin level, absolute neutrophil account) with respect to marrow adipose tissue content (p>0.05 for all values).

### Assessment of overall vascular density and CD34+ vascular morphologies

Arteriolar and sinusoidal niches have been well-described in murine models, and aging-associated alterations in densities of specific subtypes of vascular elements have also been identified.^6–10,12,15,24–29,31–34^ Therefore, we specifically examined the CD34+ vasculature in our human specimens. We used CD34 expression intensity and cytologic features to first identify endothelial cells, then trained a machine learning algorithm to generate vascular masks across all bone marrow samples, and subsequently subtyped the identified vessels by their morphologic appearance. Using parameters such as area, aspect ratio, and hull ratio, we first separated vascular elements into “large” and “small” subtypes, and then further classified small vessels as “symmetrical” or “non-symmetrical” (**Supplemental Data Fig. 5**). In each age category we observed a range of overall vascular density and relative proportions of each vessel subtype. We observed no significant difference in overall vessel density, or the densities of large or small symmetrical vessels between age groups.

Interestingly, however, we identified a conspicuous and statistically significant increase in density of small non-symmetrical vessels between the youngest and oldest marrows (0.7±0.68 mm vs. 1.1±0.95 mm, p=0.013). Recently, Sacma et al. identified a consistent overall vascular volume, and no difference in the metaphyseal sinusoidal content between young and old mice, but found a disorganized orientation of the vasculature in the epiphyses/metaphyses of aged marrows, with enrichment for vessels with decreased length and diameter.^15^ Our findings raise the possibility of a similar aging-associated remodeling of the vasculature in human bone marrows.

### Aging is associated with morphologic changes in HSPCs and megakaryocytes

Interestingly, although the proportions of several cell types (e.g. HSPCs, megakaryocytes) did not vary between age groups, our results revealed notable differences in cell sizes/morphology. We found that HSPCs (52.04±3.12 μm^2^ vs. 59.65±3.78 μm^2^, p=0.004), myeloblasts (59.43±2.98 μm^2^ vs. 69.27±3.32 μm^2^, p=0.0002), and promyelocytes (57.96±4.51 μm^2^ vs. 62.27±6.13 μm^2^, p=0.0185) were smaller in young individuals compared to individuals of both intermediate and oldest age (**Figure 3A**). HSPCs from older human individuals have been previously shown to include an increasing proportion of cells in non-G0 stages of the cell cycle.^60^ More recently, Lengefeld et al. identified an increasing proportion of larger HSPCs from older individuals, and determined that these larger cells exhibit diminished quiescence and have a reduced hematopoietic reconstitution potential.^61^ Our findings confirm that larger HSPCs comprise an increasing proportion of the total HSPC pool with aging. Conversely, megakaryocytes were much larger in younger individuals compared to both older age groups (270.42±54.16 μm^2^ vs. 178.06±58.56 μm^2^, p=0.0012), and, interestingly, megakaryocyte size showed a borderline-significant positive correlation with platelet counts (Spearman R =0.357, p=0.056) [**Figure 3B, Supplemental Data Fig. 6**]; a similar association has very recently been shown in mouse models,^62^ and therefore we hypothesize that smaller megakaryocytes less efficiently produce proplatelets and/or platelet buds, leading to reduced platelet deposition into the peripheral circulation via sinusoids. Prior studies have identified an aging-associated bias toward megakaryopoiesis in preclinical models.^63–65^ Thus, we considered that the skewing toward smaller megakaryocytes in older individuals could reflect a bias toward megakaryopoiesis in bone marrow tissues.

**Figure 3.**
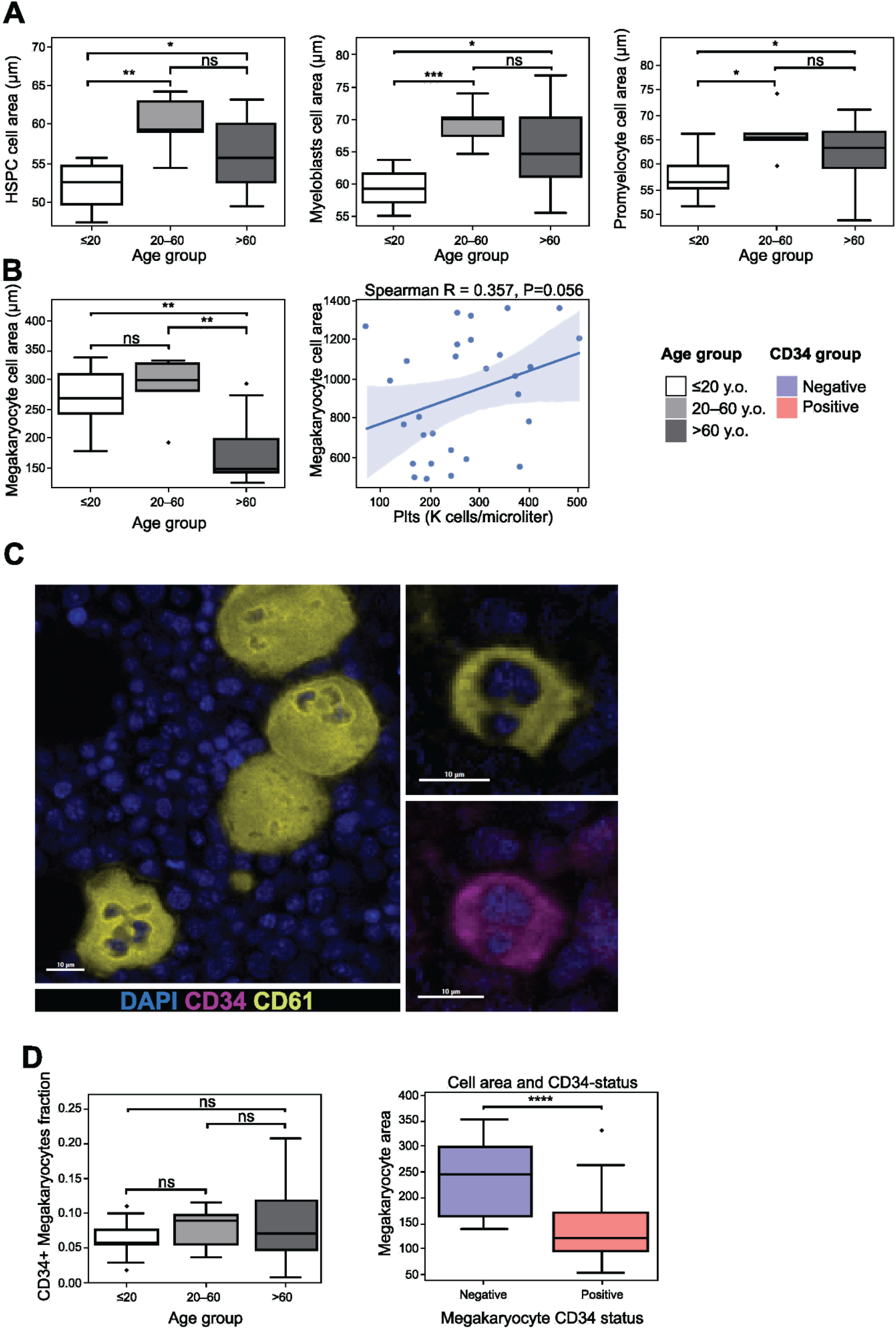
Aging-associated alterations in cell type morphologies. **A**. Pairwise comparisons between age groups for specific cell types shows increasing sizes of HSPCs (area 52.04±3.12 μm^2^ vs. area 59.65±3.78 μm^2^, p=0.004), myeloblasts (area 59.43±2.98 μm^2^ vs. area 69.27±3.32 μm^2^, p=0.0002), and promyelocytes (57.96±4.51 vs. 62.27±6.13, p=0.0185) with increasing age. **B**. Conversely, megakaryocytes were significantly larger in ≤20 year old marrows (270.42±54.17 μm^2^ vs. 178.06±58.57 μm^2^, p =0.001), and megakaryocyte size showed a borderline-significant positive correlation with platelet counts (Spearman R =0.357, p=0.056). **C**. Representative MxIF images of large mature megakaryocytes in a ≤20 year old BM (left) and a small megakaryocyte in a >60 years BM (top right). By visual assessment, a subset of small megakaryocytes exhibited cytoplasmic CD34 expression (bottom right). **D**. No significant difference between age groups in the observed proportion of CD61+ megakaryocytes that also have detectable CD34 expression (top: p>0.05 between groups); however, CD61+ megakaryocytes that also have detectable CD34 expression are significantly smaller in size (bottom: 230.7 μm^2^ vs. 120.7 μm^2^, p=0.000002).

Based on visual assessment, large mature megakaryocytes were more easily identifiable in young marrows, but in older marrows we found more conspicuous small/hypolobated forms; interestingly, these cells more frequently exhibited cytoplasmic expression of CD34, a phenotype compatible with megakaryocytic progenitors (MkPs) [**Figure 3C**]. As anticipated, CD61+ megakaryocytic elements co-expressing CD34 were significantly smaller than their CD34-negative counterparts (126.6±62.3 μm^2^ vs. 230.6±75.3 μm^2^, p=0.0000013) [**Figure 3D**]. We next determined the proportion of total CD61+ megakaryocytic elements expressing CD34 in each sample and found no clear relationship between age and proportion. These data suggest that the pool of MkPs may remain relatively constant with normal aging, but in older marrows megakaryocytes might less frequently reach a terminally-differentiated state associated with mature morphologic features.

### Spatial localization of HSPCs and other hematopoietic elements is non-random and variably consistent with aging

We next sought to specifically interrogate the topographic localization of HSPCs and determine whether well-described murine niches exist in human marrows. To address this question, we performed permutation testing (see Methods, and **Figure 4**, bottom right) for every positively identified cell type across all patient samples, including HSPCs, to interrogate for a proximity relationship with respect to megakaryocytes and structural components (e.g., bone trabeculae, fat, and CD34+ vasculature). We selected a region of 25μm (∼2-3 cell thickness) from bone trabeculae and CD34+ vasculature to account for potential direct cell-cell interactions occurring in the x- and y-planes. In addition, we also measured the observed distances between different cell types and structural components of interest. Our permutation analyses revealed heterogeneity within age groups, with both similarities and differences in cell type proximities between younger and older individuals on group-wise comparisons. Randomizing cell types of interest to other positions using all nucleated cell coordinates or only hematopoietic cell type coordinates produced similar results, as did adjusting the region of interest to 50μm for analyses focused on bone trabeculae or CD34+ vasculature.

**Figure 4.**
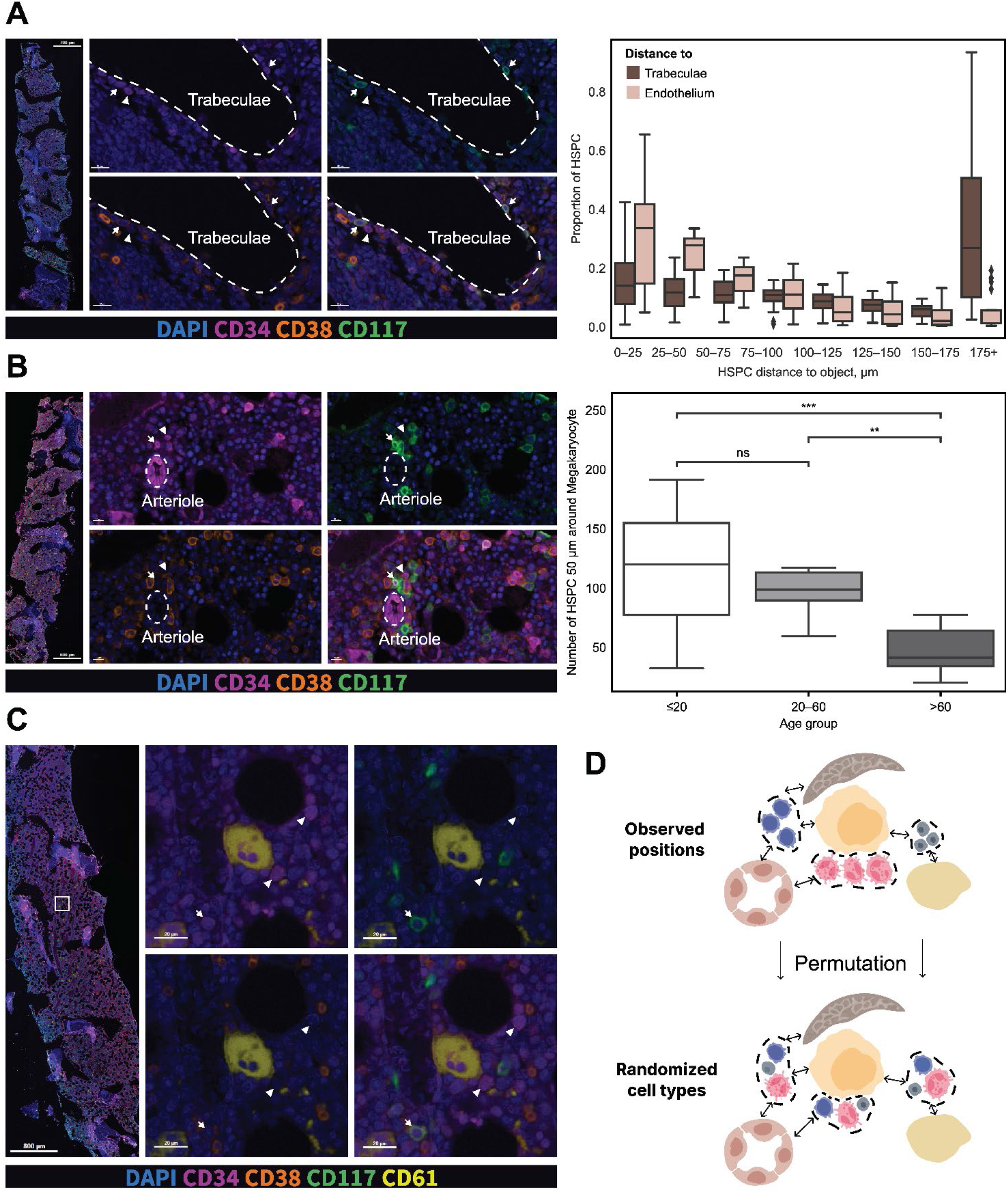
Detection of HSPCs and myeloblasts near bone trabeculae, vasculature, and megakaryocytes. **A**. Representative MxIF images from a young marrow (WCM68), including whole tissue image (left panel). Representative juxtabecular area contains HSPCs (arrowhead: CD34+CD38+/-CD117-) and myeloblasts (arrows: CD34+CD38+/-CD117+). **B**. Representative MxIF images from a young marrow (WCM66), including whole tissue image (left panel). Representative focus surrounding a CD34+ endothelium-lined arteriole contains HSPCs (arrowhead: CD34+CD38+/-CD117-) and myeloblasts (arrows: CD34+CD38+/-CD117+). *Top right*: HSPCs are located similarly close to bone trabeculae and vasculature in both young and old marrows [y-axis = HSPCs as proportion of all nucleated cells; x-axis = distance “bins” from nearest bone trabeculae or vessel]. **C**. Representative MxIF images from a young marrow (WCM60), including whole tissue image (left panel). Representative focus around a CD61+ megakaryocyte shows nearby HSPCs (arrowhead: CD34+CD38+/-CD117-) and a myeloblast in the vicinity (arrows: CD34+CD38+/-CD117+). *Right middle panel*: the absolute number of HSPCs within 50μm of CD61+ megakaryocytes borders is greater in ≤20 years marrows compared to 20-60 years and >60 years bone marrows (172.16±96.26 vs. 117.56±31.8 vs 58.54±19.1, p=0.0001). **D**. Permutation testing approach: observed locations of cell types are randomized approximately 5,000 times per each case, and then proportion of cell type(s) within region proximal to bone (25μm), vasculature (μm), or megakaryocytes (50μm) is compared to proportion in randomized location (see **Supplemental Data, Fig. 7-9**).

Based on visual inspection we observed HSPCs and myeloblasts in the region proximal to bone trabeculae (**Figure 4A**). When we quantified the proportion of HSPCs within discrete distance “bins” we found a diminution in from proximal to distal to trabeculae in both ≤20 years and >60 years marrows. Based on permutation testing, we found HSPCs, myeloblasts, and promyelocytes were significantly enriched near bone trabeculae in young individuals; we observed a similar pattern in older adults but identified only a trend toward significance for myeloblasts (**Supplemental Data Fig. 7**). These findings indicate that the immediate juxta-trabecular area represents at least one concentrated focus of early myelopoiesis in humans, and that this spatial anatomy remains relatively constant with aging. In older individuals, we also found a relative enrichment of plasma cells and CD34+ endothelial cells near bone trabeculae, suggesting a remodeling of the endosteal area with aging. NHEs were similarly enriched near bone at the extremes of age; we hypothesize that these cells include a heterogeneous composition of mature immune and stromal elements (e.g. osteoblasts, perivascular cells). Conversely, more differentiated cell types including MMCs, all stages of erythroid maturation, and megakaryocytes were similarly and significantly distant from bone trabeculae in almost all samples, consistent with the commonly observed distribution of these cell types on conventional histomorphologic evaluation.

Based on visual inspection we also observed HSPCs and myeloblasts in the region proximal to CD34+ vascular elements (**Figure 4B**). When we quantified the proportion of HSPCs within discrete distance “bins” we found a diminution in from proximal to distal to vasculature in both ≤20 years and >60 years marrows. Using permutation testing, we found that HSPCs were similarly enriched near vasculature at the extremes of age (**Supplemental Data Fig. 8**). While myeloblasts were also enriched in this region in young individuals, we observed only a trend toward enrichment in older individuals. Similarly, we found trends toward enrichment in both age groups for promyelocytes. Interestingly, we observed an enrichment for megakaryocytes near CD34+ vasculature in older individuals, but a conspicuous “distant” pattern in young individuals. Our permutation testing approach hinges on the use of cell centroids (i.e., nuclei) for measurement, and young marrows containing mostly large mature megakaryocytes. Thus, we hypothesize that this finding may be a result of the relatively larger distance between CD34+ vasculature and cell centroids (nuclei) in larger megakaryocytes, even in instances of close proximity based on visual inspection. In contrast to the region proximal to bone trabeculae, we found non-hematopoietic elements to be excluded from the perivascular zone.

### Evidence for a potential megakaryocytic niche in humans

A few prior studies have demonstrated the existence of a quiescent megakaryocytic niche for HSPCs in preclinical models.^36,37^ Our evaluation of cell type proportions and morphologies (cell areas) in human tissues revealed no significant differences in HSPC percentage between age groups, but rather we found the presence of significantly smaller megakaryocytes with increasing age, which led us to inspect for a spatial relationship between HSPCs and megakaryocytes in human marrow. Permutation testing revealed a non-significant trend toward a spatial relationship in younger individuals only, in spite of the larger distance between the centroids (nuclei) of HSPCs and larger megakaryocytes found in young marrows; we found no such relationship in older individuals (**Supplemental Data Fig. 9)**. On visual assessment, we observed HSPCs and myeloblasts localized near megakaryocytes, particularly in young bone marrows (**Figure 4C**). To account for the differences in megakaryocyte sizes in different age groups, and the methodology for cell-cell distance measurements (i.e., centroid/nucleus to centroid/nucleus) we used a dynamic range for each megakaryocyte (megakaryocyte size + 50 μm). The absolute number of HSPCs within 50 μm of CD61+ megakaryocytes borders was greater in marrows from ≤20 years individuals compared to both 20-60 years and >60 years age groups (172.16±96.26 vs. 117.56±31.8 vs 58.54±19.1, p=0.0001). These findings raise the possibility of an active megakaryocytic niche in human tissues, which may become less prominent upon aging.

### Detection of unique cell communities

In order to dissect subtle features of the marrow topography undetectable by light microscopy and other conventional methods, we employed a cell communities analysis at whole tissue scale (see Methods).^66^ By combining cellular and structural features in our model, we identified 6 unique communities, which varied in their compositions of structural elements (e.g. bone trabeculae, CD34+ vasculature, and fat) and cellular constituents (**Figure 5A**). To confirm the accuracy of community detection we performed histopathologic re-examination of digitized H&E-stained sections alongside their MxIF preparations, and corresponding Voronoi plots (cell maps) from select cases. Clusters differed with regard to their composition by both structural components and individual cell types (**Figure 5A**). MMCs and erythroid normoblasts were relatively prominent in clusters 0 and 2, respectively, while cluster 1 was notably enriched in non-hematopoietic elements. Interestingly, HSPCs were equally frequent in clusters 1 and 5 (p>0.05) [**Figure 5B**]; however, cluster 1 included a relative diminution of promyelocytes and proerythroblasts, suggesting decreased HSPC lineage output in these areas. In pairwise analyses comparing individuals ≤20 years and >60 years, we observed statistically significant reductions in the proportions of clusters 0 (0.232±0.157 vs. 0.136±0.155, p=0.0038) and 3 (0.265±0.07 vs. 0.175±0.068, p=0.004).

**Figure 5.**
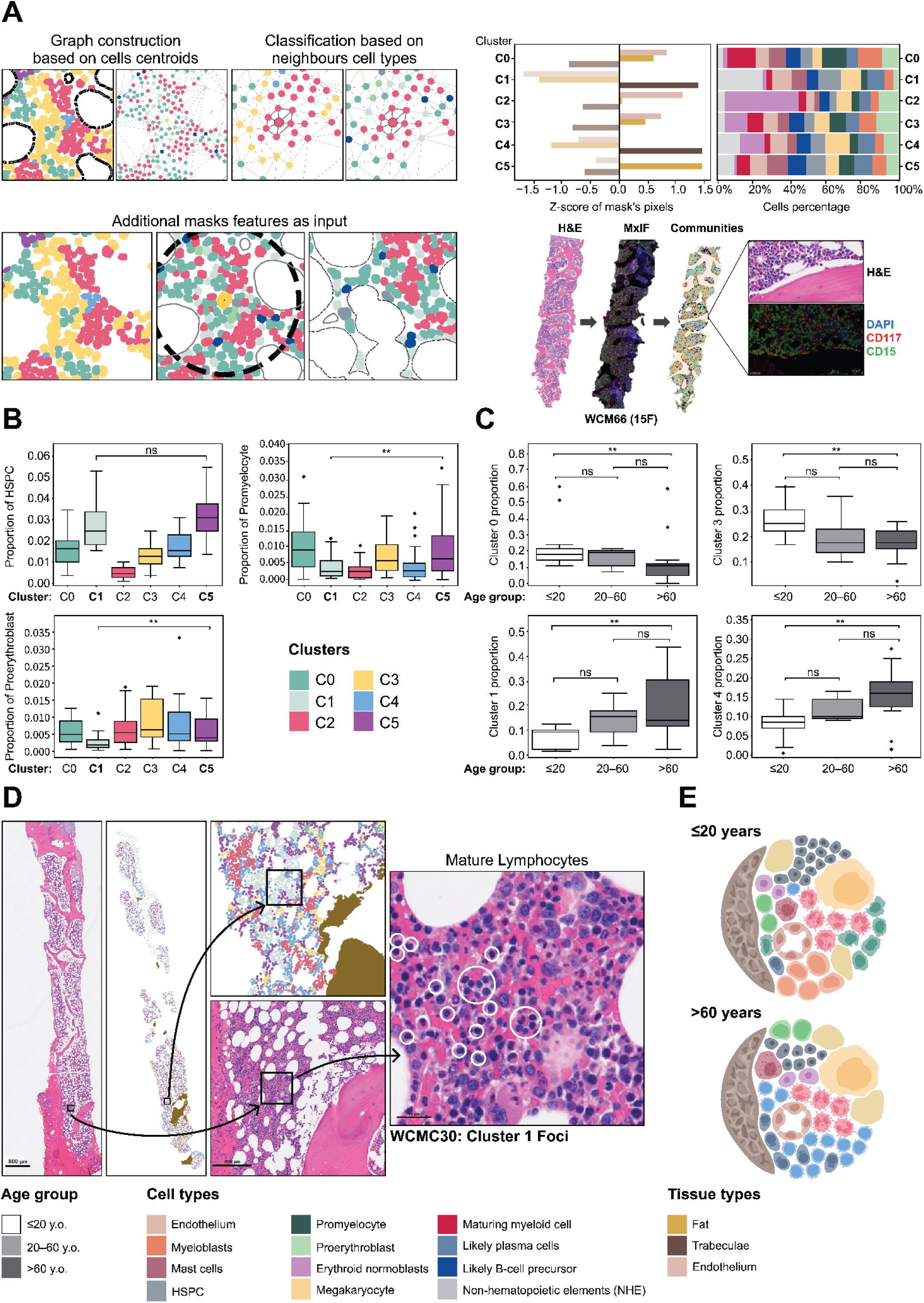
Detection of unique cell communities and evidence of inflammaging. **A**. A 75μm region was utilized for community detection. Six unique cellular/structural communities were detected, differentiated by proportions of constituent hematopoietic and non-hematopoietic elements, and relative contributions from bone trabeculae, CD34+ vasculature, and fat (left). Schematic of paired hematoxylin and eosin-stained section, MxIF preparation, and topographic communities map (case WCM66), with a highlighted juxtatrabecular focus enriched for immature myeloid elements including myeloblasts and promyelocytes, with distal progression toward MMCs. **B**. HSPCs are relatively enriched in clusters 1 (0.0282±0.1 vs 0.0178±0.006) and 5 (0.0315±0.095 vs 0.0178±0.006) vs third most abundant cluster 4. Cluster 1 exhibits a relative depletion of promyelocytes (0.0038±0.0036 vs. 0.0098±0.009, p=0.0015) and proerythroblasts (0.0026±0.0023 vs. 0.0059±0.0045, p=0.0005) compared to Cluster 5. **C**. Clusters 0 (0.232±0.157 vs. 0.136±0.155, p=0.0035) and 3 (0.265±0.071 vs. 0.175±0.068, p=0.0034) are relatively enriched in ≤20 years marrows as compared to >60 years marrows. Clusters 1 (0.067±0.043 vs. 0.187±0.123, p=0.0015) and 4 (0.085±0.043 vs. 0.161±0.072, p=0.0025) are relatively enriched in >60 years marrows as compared to ≤20 years marrows. **D**. Mapping of Cluster 1 foci back to H&E-stained WSI reveals conspicuous mature lymphocytes by histomorphologic assessment. **E**. Hypothetical model of increased immune cells comprising the NHEs (light blue cells) enriched in Cluster 1 compatible with “inflammaging”. *0.01 < p <= 0.05, **0.001 < p <= 0.01, ***0.0001 < p <= 0.001, ****p<0.0001.

Conversely, individuals >60 years displayed greater proportions of clusters 1 (0.067±0.043 vs. 0.187±0.122, p=0.0015) and 4 (0.085±0.043 vs. 0.161±0.072, p=0.0027) [**Figure 5C**]. Although our MxIF assay design precluded the ability to definitively classify non-hematopoietic elements by phenotype, we performed a histologic examination of representative tissue areas comprised by cluster 1 in individuals >60 years (e.g. WCM30, WCM35, WCM53, WCM56), and identified conspicuous cells with cytomorphologic features compatible with mature lymphocytes (**Figure 5D, Supplemental Data Figure 10**).

Thus, we hypothesize that the proportional increase in the percentage of cluster 1, and to a lesser extent cluster 4, observed in older individuals may at least partially represent “inflammaging”,^67^ a well-described phenomenon linked to chronic inflammation that can contribute to cytopenias, diminished immune responses, and an increased risk of neoplasia (**Figure 5E**). Furthermore, the diminished lineage output from HSPCs in these cluster 1 areas may be associated with the presence of low-grade chronic inflammation.

## DISCUSSION

Our existing understanding of the bone marrow spatial cellular architecture and microenvironment is a result of extensive work by many groups, primarily using murine model systems. Although these studies have been critical in identifying and dissecting components of the HSPC niche, the generalizability of the resulting knowledge, particularly with respect to the human bone marrow, has remained largely unknown. For many years, technical challenges associated with imaging human bone marrow tissues at the level of detail required to dissect the niche have precluded in-human corroboration of findings from murine modeling. However, with recent advances in imaging techniques and computational bioinformatics these challenges can now be overcome.^54,58^ Here, we have described a pipeline for single-cell resolution-based spatial mapping of the human bone marrow using decalcified, fixed, and paraffinized core biopsy tissues originally procured for routine clinical diagnostics. Importantly, this pipeline can be applied to serial samples collected from individuals, creating an opportunity to study the longitudinal evolution of a variety of bone marrow pathologies with single-cell level resolution in the context of preserved marrow architecture.

Using multiparametric in situ imaging and AI-driven analytical techniques, we demonstrate aging-associated alterations in human hematopoiesis, at both the level of the single cell and with respect to regional anatomy, including marrow adiposity and vascular morphology. Although we found that the frequencies of various cell types within the hematopoietic compartment did not differ significantly with aging specifically, we observed heterogeneous adiposity within age groups, and an interesting association between higher fat content and constrained HSPC differentiation toward granulocytic and erythroid progenitors, as previously reported in mice.^59^ Subsequent assessment of the morphological features of individual cell types revealed an association between older age and the presence of larger HSPCs, but smaller megakaryocytes. Lengefeld et al. recently described a close relationship between increasing HSC age, cell size, and diminished quiescence;^61^ some of their findings confirm work from prior studies using human-derived and sorted HSCs.^60^

The spatial positioning of HSPCs near bone and vasculature has been described in preclinical models by many groups in recent decades; our data confirmed this localization in human marrow, but did not reveal significant changes associated with aging. Our permutation testing approach also enabled us to assess for enrichment or relative exclusion of many other cell types within these regions of the marrow; based on these data we conclude that maturing myelopoiesis is predominantly initiated in the endosteal region, with more differentiated elements primarily localized to interstitial areas of the marrow. Similarly, megakaryocytes and almost all phases of erythroid maturation are located distal to bone and vasculature in both young and old individuals. Given that evidence of a megakaryocytic niche has been elegantly shown in murine models,^36,37^ we investigated the spatial relationship between megakaryocytes and HSPCs in human marrow. Remarkably, we found that HSPCs were relatively enriched within the vicinity of megakaryocytes to a greater extent in younger versus older marrows, providing evidence of a potential megakaryocytic niche in humans, the activity of which may diminish with aging. Overall, these findings confirm that the spatial anatomy of human bone marrow is non-random and suggest that particular regions of the marrow exhibit localized, aging-associated compositional alterations.

A particularly important facet of our study is the detection of cell communities, which allowed us to dissect the microarchitecture and spatial arrangement of numerous cell types at a resolution far exceeding that of light microscopy or conventional visual assessment of chromogenic immunohistochemistry. Using this approach, we found that HSPCs in younger marrows are more frequently present in communities enriched for their progeny; in older individuals, we find HSPCs more often associated with non-hematopoietic elements. Guided by digital cell community maps, we focused a histologic evaluation on tissue areas enriched for these non-hematopoietic elements (e.g. Cluster 1 foci). Based on this analysis, we observed conspicuous mature lymphocytes in these areas, potentially representing a component of “inflammaging”;^67^ further phenotypic and functional characterization of these non-hematopoietic elements is a key focus of ongoing investigation.

We demonstrate the capability to perform MxIF staining, whole tissue imaging, and AI/deep learning-based analytics to dissect the spatial biology of the human marrow at single cell resolution using readily-available archival specimens; importantly, the staining and imaging workflow we describe is widely accessible and lacks significant barriers for implementation. One practical limitation of this study relates to the indications for why bone marrow specimens were originally collected from the included individuals (e.g., staging procedure for another malignancy, or very mild cytopenias); although we cannot account for potentially confounding factors, all bone marrow specimens were considered normal by comprehensive diagnostic evaluation. Nonetheless, future studies may benefit from use of bone marrow specimens from volunteers. Current limitations of our analysis pipeline include a relatively limited number of antigens profiled concurrently, and a restriction to two-dimensional tissue imaging; although the output data may be descriptive, assessments of longitudinal samples collected from individual patients will provide the opportunity to study the intact marrow topography and entire microenvironment along the course of disease evolution. Moreover, this type of approach will be further enhanced in the years to come by parallel technological advances that will enable the detection of many more biomarkers in single tissue sections. Thus, our findings pave the way for future spatial and temporal interrogation of the human bone marrow microenvironment, in a variety of physiologic and pathologic states, including metastatic disease and numerous hematologic malignancies such as myelodysplasia, acute leukemia, myeloma, and other marrow-based myeloid and lymphoid neoplasia.

## METHODS

### Case Selection

A total of 29 individuals were included in the study. Decalcified, bouin-fixed, and paraffin-embedded iliac crest bone marrow (BM) biopsy material was selected for the study and de-identified prior to testing, upon approval by the institutional review board (IRB) and applicable committees at Weill Cornell Medical College/NewYork-Presbyterian Hospital (WCM/NYP). Per standard clinical processing protocols established at WCM/NYP, archival BMs were fixed in Bouin solution (American MasterTech, Lodi, CA, USA) and decalcified (Epredia Decalcifying Solution, Fisher Scientific, Waltham, MA) before routine processing and paraffin embedding. Whole BMs (N=29) were evaluated originating from individuals who had no evidence of a bone marrow disease based upon routine comprehensive diagnostic evaluation (“NBM”; <20 years = 11, 20-60 years = 5, >60 years = 13). Hematoxylin and eosin (H&E)-stained preparations from each candidate sample were re-reviewed (SSP) to ensure sufficient quantity and quality of material to be utilized for subsequent testing. Whole slide images (WSIs) of each H&E-stained preparation were captured using the Vectra Polaris Automated Quantitative Pathology Imaging System (Akoya Biosciences, Marlborough, MA).

### Clinicopathologic and Laboratory Data

The electronic medical record (EMR) for each patient included in the study was reviewed to obtain data concurrent with the evaluated biopsy specimen, including time of collection, patient body mass index (BMI), and complete blood count (CBC) data including hemoglobin concentration (Hb), mean corpuscular value (MCV), absolute neutrophil count (ANC), and platelet count (Plt). All diagnostic hematopathology reports were re-reviewed (SSP, JTG) to confirm the original diagnosis of a normal bone marrow, and extract data related to manual aspirate differential counting, including myeloid to erythroid (M:E) ratios, and percentages of blasts, promyelocyctes, proerythroblasts, granulocytes, monocytes, erythroid normoblasts, and plasma cells.

### Multiplex Immunofluorescence Tissue Staining

Multiplexed immunofluorescence (MxIF) was performed using the Opal system (Akoya Biosciences, Marlborough, MA) by staining 4 micron-thick Bouin-fixed, paraffin-embedded whole-tissue sections from decalcified human BM core biopsy specimens in a Bond RX automated tissue stainer (Leica Biosystems, Buffalo Grove, IL), as described previously ^80,81,87,88^. Briefly, tissue sections were first deparaffinized prior to EDTA-based antigen retrieval (Leica ER2 solution, 20 minutes). A cyclical staining protocol was then performed, with horseradish peroxidase-mediated deposition of tyramide-Opal fluorophore constructs (Akoya Biosciences, Marlborough, MA) in each cycle, with intervening application of heat, citrate-based epitope retrieval solution (Leica ER1), and Bond Wash Solution (Leica) to execute stripping of primary/secondary antibody complexes between staining cycles. Finally, 4’, 6-diamidino-2-phenylindole (Spectral DAPI, Akoya Biosciences) was applied per provided protocols to label nuclei. Antibodies were selected from a menu of extensively validated and clinically tested clones in our CLIA laboratory (Weill Cornell Medicine/NewYork-Presbyterian Hospital, New York, NY). The following panel of primary antibody/fluorophore pairs was applied to all cases, in a sequential order as shown: 1) Opal 480/CD71 (1:80, 10F11, Leica concentrate), 2) Opal 520/CD61 (Ready-to-use, 2F2, Leica), 3) Opal 570/anti-CD117 (1:100, D3W6Y, Cell Signaling), 4) Opal 620/CD38 (1:50, 38C03(SPC32), Invitrogen), 5) Opal 690/anti-CD34 (1:100, QBEND/10, Invitrogen), 6) Opal 780/CD15 (1:50, MMA, BD Biosciences). All slides were coverslipped using ProLong™ Diamond Antifade Mountant (Invitrogen). WSI scans were subsequently obtained at 20X magnification using the Vectra Polaris Automated Quantitative Pathology Imaging System (Akoya Biosciences). WSIs were tiled in Phenochart (v1.1, Akoya Biosciences); image tiles were subsequently spectrally unmixed in InForm (v2.4.8, Akoya Biosciences). Unmixed tiles were finally fused together in HALO (v3.3.2541.231, Indica Labs, Albuquerque, NM) to generate a single multi-layered TIFF image file for each sample, which was used in downstream analyses. Spectrally-unmixed WSIs were parsed through custom artificial intelligence/machine learning-based Python pipelines developed and applied by BostonGene. Representative images presented in figures were generated using Phenochart.

### Cell Segmentation

A convolutional neural network (CNN) was trained to perform cell segmentation. Mask-RCNN architecture with a ResNet-18 backbone was chosen to detect cells as separate instances. The model was trained on manually annotated images. The training set consisted of 780 images of different types of tissue (kidney, tonsil, lung, colon) separately stained with the CO-Detection by indEXing (CODEX) platform and 156 images of human bone marrow tissues stained and imaged as above (see ‘Multiplex Immunofluorescence Tissue Staining’). The validation set consisted of 20 images of BM only. All annotated images had side size equal to 256 pixels.

The model was designed to use a four-channel image as input. The first image corresponded to nuclei stained with DAPI, the second image corresponded to the most common membrane marker (NaKATPase for images stained with CODEX and CD71 for bone marrow images), and the third image corresponded to a stack of other membrane markers. The fourth channel was required for adequate detection of megakaryocytes and consisted of an empty channel for images imaged with CODEX and a CD61 channel for BM images.

For training and inference purposes all images were normalized to a 0-1 range.

### Structure Masks Generation

Fat and bone trabeculae masks were generated by a CNN with UNet++ architecture and efficient net-b1 encoder. The model required all available markers in the bone marrow staining panel as inputs (DAPI, CD15, CD71, CD117, CD34, CD38, CD61) and provided fat and bone trabeculae masks as outputs. The dataset consisted of >16,500 images with sizes of training and validation sets of 75% and 25%, respectively. Fixed training and testing image distribution were used during network training.

In total two models were trained. A pathologist then selected the best mask for every case and every mask type. For cases where both trained models failed a pathologist performed manual correction (7 manually generated masks for trabeculae in total).

For training and inference purposes all images were normalized to a 0-1 range and then z-score normalization was applied.

Endothelium masks were generated by a CNN with DeepLabV3+ architecture and efficient net-b0 encoder. A single channel (CD34 marker) was used as the input and a single channel endothelium mask was generated as the output. The dataset consisted of 40 annotated images with dimensions of 1024×1024 pixels. To expand the total training set size the original images were split into tiles of 256×256 pixels dimension with intersection of 128 pixels between tiles. For the validation set the intersection was set to 0. This approach resulted in 1862 training images and 32 validation images. For training and inference purposes all images were normalized to 0-1 range.

### Classification of CD34+ Endothelium-Lined Vessels

Separate instances of blood vessels were obtained from endothelium masks, and further sub-classified based on geometric parameters including hull ratio, aspect ratio and area. “Small symmetrical” vessels were defined using hull ratio >0.8, aspect ratio >0.5, and area <400 μm^2^. “Small non-symmetrical” vessels were defined using hull ratio >0.5, aspect ratio >0.3, and area <310 μm^2^. Parameters were adjusted manually to account for additional variations in vessel morphology.

To separate large vessels (likely sinusoids) we used the object contours descriptors. For each vessel object we calculated the perimeter, area, major and minor axis length and several other measurements. Based on it we performed the clusterization into known types of vasculature.

### Cell Typing

Cell typing was performed based on the probability of signal. To obtain these probabilities a classification convolutional network was trained to predict presence or absence of marker signal along cell contours. The model was trained with a three-channel input image: DAPI (channel 1), marker of interest (channel 2), segmentation mask from segment of interest (channel 3). All three channels were centered around the cell of interest and normalized to a 0-1 range. The image size was set to 128×128 pixels. The model’s output was a softmax on two classes: presence and absence of the signal on a given cell, enabling natural normalization to a 0-1 range.

A ResNet-50 architecture was used for the CNN. The dataset consisted of 890 training images and 650 validation images, with a fixed split between training and validation runs.

Cell type classification was performed after signal probabilities were obtained. For each cell type we first defined a vector of the “ground truth” (or signature) expression. For example, the CD71 vector position for cells typed as “erythroid normoblasts” was set to one, and zero for all other cell types. After all vectors were defined the final prediction was generated as a closest match between the predicted probabilities vector and the reference “ground truth” vector by a cosine similarity.

To separate populations of endothelial cells from HSPCs, blood vessel masks were used. CD34+ cells with greater than 50% of their cell area intersecting a blood vessel mask were typed as endothelial cells, while all others were subsequently considered as hematopoietic cells expressing CD34. Cell area was used to distinguish likely B-cell precursors (hematogones) from likely plasma cells, using a threshold value of 87.5. Larger cells were classified as plasma cells, and others as likely B-cells. Additional details are provided in Supplemental Methods.

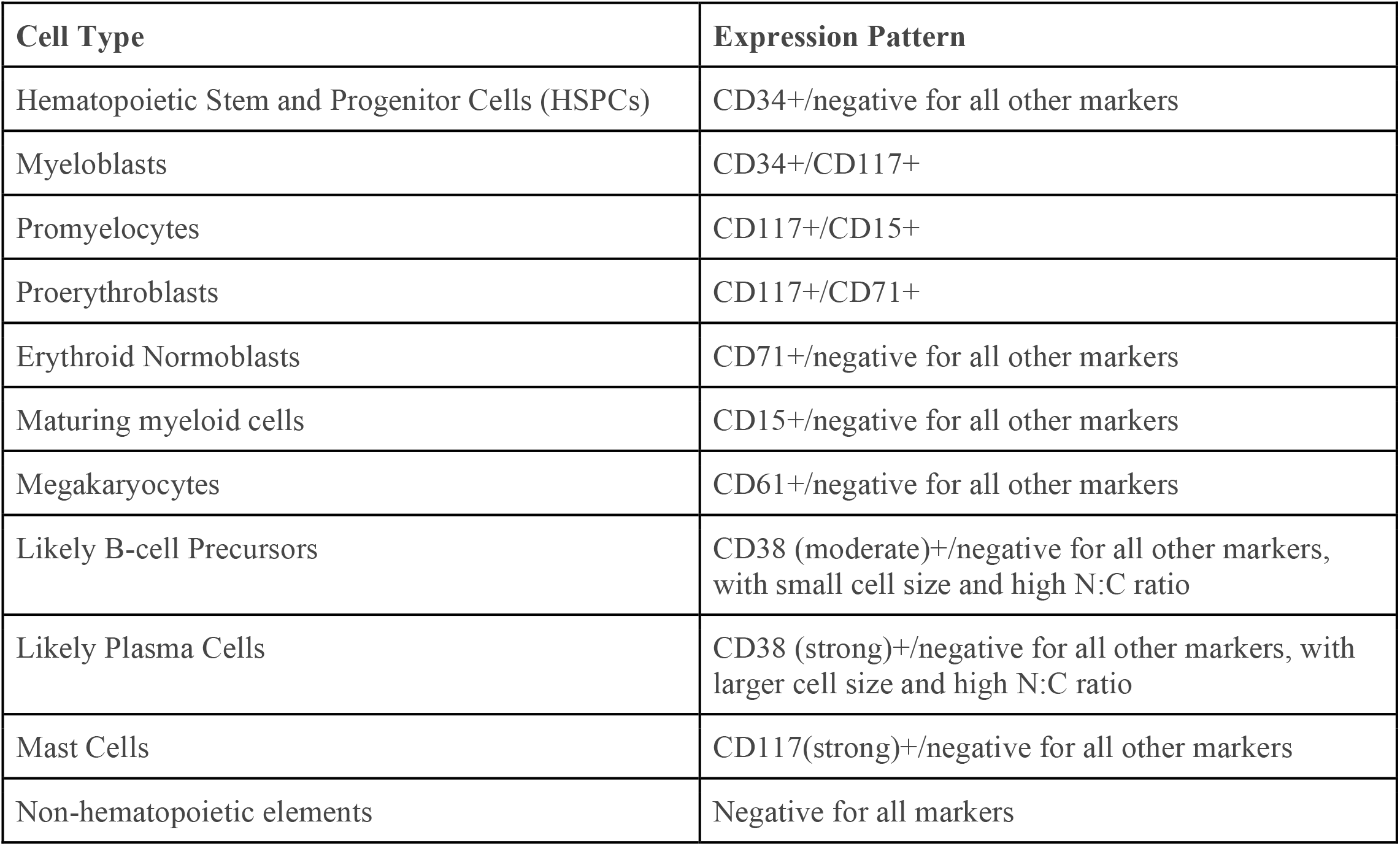

### Permutation Testing

We used a permutation test to interrogate for enrichment or avoidance of cell types of interest with respect to topographical structures such as fat, bone trabeculae, vessels, and additionally for relationship with megakaryocytes. A 25 μm and 50 μm radii from objects of interest were tested for each replica of a permutation. Coordinates for all nucleated cells (DAPI+), or only x-y coordinates for hematopoietic cells (i.e. marker positive) were separately tested as locations for permuted cells. To obtain a p-value of significance we applied 5000 random permutations per patient, and compared original (observed) distances of a given cell type to the object of interest with distances generated following random shuffling. Pearson’s method was used to combine p-values of individual samples for established age groups.

### Community Analysis

Community analysis was performed based on a graph of cell centroids. The graph was constructed by a Delaunay triangulation algorithm with additional removal of edges with length exceeding 100 *μ*m. In a resulting graph each node additionally contained encoded information about its cell type, mean distance to its neighbors and percentages of structure masks (trabeculae, fat, endothelium) within a 75 *μ*m radius underneath node center. This dimension was selected to account for the nearest neighbors and structures of each cell centroid, within approximately 3 cell diameters away; this region size has also been previously utilized and validated for similar community analyses.^68^

Adversarially Regularized Variational Graph Auto-Encoder^69^ was then trained for 100 epochs in an unsupervised manner to obtain representative embedding for each node. The model consisted of encoder and decoder parts. The encoder produced vectors with length of 32 which were further used as encoded neighborhood features for clusterization.

KMeans clusterization into 6 clusters was then performed on obtained feature vectors. The number of clusters was optimized by minimization of Calinski-Harabasz index and maximization of Silhouette and Davies-Bouldin scores. Additional details are provided in Supplemental Methods. Community clusters were classified by dominant cell type and mean percentage composition by structural masks. Detailed community cluster descriptions are provided in the following table.

| Community Label | Description |
| --- | --- |
| Cluster 0 | Maturing myeloid cells + Myeloblasts + Promyelocytes |
| Cluster 1 | Non-hematopoietic elements (Likely Lymphocytes) |
| Cluster 2 | Erythroid Normoblasts |
| Cluster 3 | Erythroid Normoblasts + Maturing myeloid cells + Proerythroblasts |
| Cluster 4 | Erythroid Normoblasts + Non-hematopoietic elements |
| Cluster 5 | Maturing myeloid cells + HSPCs + Non-hematopoietic elements |

### Statistical Analyses

Statistical analyses were performed in Python (v3.10), using the scipy (v1.7.3) and statsmodels (0.13.1) packages. The Mann-Whitney U test was used for group-wise comparisons and Spearman’s correlation coefficient was used for correlation testing, if not otherwise specified. The Benjamini-Hochberg procedure was used to correct for testing of multiple comparisons. Statistical significance was set to p<0.05, unless otherwise specified.

## DATA AVAILABILITY

Requests for raw data can be submitted in writing via e-mail to the corresponding author.

## AUTHOR CONTRIBUTIONS

Experiments were designed, executed and/or analyzed by S.S.P., A.S., V.S., A.V., I.G., and D.T. I.V., C.U. and T.P. performed multiplex tissue staining and imaging. The manuscript was written by S.S.P. with input from all authors. S.S.P. conceived of and designed the study, and supervised the work.

## ACKNOWLEDGEMENTS

This work was supported by the Department of Pathology and Laboratory Medicine, Weill Cornell Medical College (start-up funding to S.S.P.). Tissue staining and imaging was performed in the Multiparametric In Situ Imaging (MISI) Laboratory of the Department of Pathology and Laboratory Medicine, Weill Cornell Medical College. The authors thank Dr. Daniel Lucas (Cincinnati Children’s Hospital) and Dr. Ruben Carrasco (Dana-Farber Cancer Institute) for critical reading and input on the manuscript.

## REFERENCES

1. Jagannathan-Bogdan M, Zon LI. Hematopoiesis. Dev. Camb. Engl. 2013;140(12):2463–2467.

2. Pinho S, Frenette PS. Haematopoietic stem cell activity and interactions with the niche. Nat. Rev. Mol. Cell Biol. 2019;20(5):303–320.

3. Crane GM, Jeffery E, Morrison SJ. Adult haematopoietic stem cell niches. Nat. Rev. Immunol. 2017;17(9):573–590.

4. Lo Celso C, Fleming HE, Wu JW, et al. Live-animal tracking of individual haematopoietic stem/progenitor cells in their niche. Nature. 2009;457(7225):92–96.

5. Xie Y, Yin T, Wiegraebe W, et al. Detection of functional haematopoietic stem cell niche using real-time imaging. Nature. 2009;457(7225):97–101.

6. Méndez-Ferrer S, Michurina TV, Ferraro F, et al. Mesenchymal and haematopoietic stem cells form a unique bone marrow niche. Nature. 2010;466(7308):829–834.

7. Kunisaki Y, Bruns I, Scheiermann C, et al. Arteriolar niches maintain haematopoietic stem cell quiescence. Nature. 2013;502(7473):637–643.

8. Nombela-Arrieta C, Pivarnik G, Winkel B, et al. Quantitative imaging of haematopoietic stem and progenitor cell localization and hypoxic status in the bone marrow microenvironment. Nat. Cell Biol. 2013;15(5):533–543.

9. Spencer JA, Ferraro F, Roussakis E, et al. Direct measurement of local oxygen concentration in the bone marrow of live animals. Nature. 2014;508(7495):269–273.

10. Acar M, Kocherlakota KS, Murphy MM, et al. Deep imaging of bone marrow shows non-dividing stem cells are mainly perisinusoidal. Nature. 2015;526(7571):126–130.

11. Hawkins ED, Duarte D, Akinduro O, et al. T-cell acute leukaemia exhibits dynamic interactions with bone marrow microenvironments. Nature. 2016;538(7626):518–522.

12. Itkin T, Gur-Cohen S, Spencer JA, et al. Distinct bone marrow blood vessels differentially regulate haematopoiesis. Nature. 2016;532(7599):323–328.

13. Duarte D, Hawkins ED, Akinduro O, et al. Inhibition of Endosteal Vascular Niche Remodeling Rescues Hematopoietic Stem Cell Loss in AML. Cell Stem Cell. 2018;22(1):64–77.e6.

14. Pinho S, Marchand T, Yang E, et al. Lineage-Biased Hematopoietic Stem Cells Are Regulated by Distinct Niches. Dev. Cell. 2018;44(5):634–641.e4.

15. Saçma M, Pospiech J, Bogeska R, et al. Haematopoietic stem cells in perisinusoidal niches are protected from ageing. Nat. Cell Biol. 2019;21(11):1309–1320.

16. Kokkaliaris KD, Kunz L, Cabezas-Wallscheid N, et al. Adult blood stem cell localization reflects the abundance of reported bone marrow niche cell types and their combinations. Blood. 2020;136(20):2296–2307.

17. Nilsson SK, Dooner MS, Tiarks CY, Weier HU, Quesenberry PJ. Potential and distribution of transplanted hematopoietic stem cells in a nonablated mouse model. Blood. 1997;89(11):4013–4020.

18. Calvi LM, Adams GB, Weibrecht KW, et al. Osteoblastic cells regulate the haematopoietic stem cell niche. Nature. 2003;425(6960):841–846.

19. Zhang J, Niu C, Ye L, et al. Identification of the haematopoietic stem cell niche and control of the niche size. Nature. 2003;425(6960):836–841.

20. Nilsson SK, Johnston HM, Whitty GA, et al. Osteopontin, a key component of the hematopoietic stem cell niche and regulator of primitive hematopoietic progenitor cells. Blood. 2005;106(4):1232–1239.

21. Adams GB, Chabner KT, Alley IR, et al. Stem cell engraftment at the endosteal niche is specified by the calcium-sensing receptor. Nature. 2006;439(7076):599–603.

22. Zhu J, Garrett R, Jung Y, et al. Osteoblasts support B-lymphocyte commitment and differentiation from hematopoietic stem cells. Blood. 2007;109(9):3706–3712.

23. Zhao M, Tao F, Venkatraman A, et al. N-Cadherin-Expressing Bone and Marrow Stromal Progenitor Cells Maintain Reserve Hematopoietic Stem Cells. Cell Rep. 2019;26(3):652–669.e6.

24. Kiel MJ, Yilmaz OH, Iwashita T, et al. SLAM family receptors distinguish hematopoietic stem and progenitor cells and reveal endothelial niches for stem cells. Cell. 2005;121(7):1109–1121.

25. Sugiyama T, Kohara H, Noda M, Nagasawa T. Maintenance of the hematopoietic stem cell pool by CXCL12-CXCR4 chemokine signaling in bone marrow stromal cell niches. Immunity. 2006;25(6):977–988.

26. Sacchetti B, Funari A, Michienzi S, et al. Self-renewing osteoprogenitors in bone marrow sinusoids can organize a hematopoietic microenvironment. Cell. 2007;131(2):324–336.

27. Ding L, Saunders TL, Enikolopov G, Morrison SJ. Endothelial and perivascular cells maintain haematopoietic stem cells. Nature. 2012;481(7382):457–462.

28. Ding L, Morrison SJ. Haematopoietic stem cells and early lymphoid progenitors occupy distinct bone marrow niches. Nature. 2013;495(7440):231–235.

29. Greenbaum A, Hsu Y-MS, Day RB, et al. CXCL12 in early mesenchymal progenitors is required for haematopoietic stem-cell maintenance. Nature. 2013;495(7440):227–230.

30. Chen JY, Miyanishi M, Wang SK, et al. Hoxb5 marks long-term haematopoietic stem cells and reveals a homogenous perivascular niche. Nature. 2016;530(7589):223–227.

31. Kusumbe AP, Ramasamy SK, Itkin T, et al. Age-dependent modulation of vascular niches for haematopoietic stem cells. Nature. 2016;532(7599):380–384.

32. Asada N, Kunisaki Y, Pierce H, et al. Differential cytokine contributions of perivascular haematopoietic stem cell niches. Nat. Cell Biol. 2017;19(3):214–223.

33. Comazzetto S, Murphy MM, Berto S, et al. Restricted Hematopoietic Progenitors and Erythropoiesis Require SCF from Leptin Receptor+ Niche Cells in the Bone Marrow. Cell Stem Cell. 2019;24(3):477–486.e6.

34. Butler JM, Nolan DJ, Vertes EL, et al. Endothelial cells are essential for the self-renewal and repopulation of Notch-dependent hematopoietic stem cells. Cell Stem Cell. 2010;6(3):251–264.

35. Yamazaki S, Iwama A, Takayanagi S, et al. TGF-beta as a candidate bone marrow niche signal to induce hematopoietic stem cell hibernation. Blood. 2009;113(6):1250–1256.

36. Bruns I, Lucas D, Pinho S, et al. Megakaryocytes regulate hematopoietic stem cell quiescence through CXCL4 secretion. Nat. Med. 2014;20(11):1315–1320.

37. Zhao M, Perry JM, Marshall H, et al. Megakaryocytes maintain homeostatic quiescence and promote post-injury regeneration of hematopoietic stem cells. Nat. Med. 2014;20(11):1321–1326.

38. Heazlewood SY, Neaves RJ, Williams B, et al. Megakaryocytes co-localise with hemopoietic stem cells and release cytokines that up-regulate stem cell proliferation. Stem Cell Res. 2013;11(2):782–792.

39. Ludin A, Itkin T, Gur-Cohen S, et al. Monocytes-macrophages that express α-smooth muscle actin preserve primitive hematopoietic cells in the bone marrow. Nat. Immunol. 2012;13(11):1072–1082.

40. Chow A, Lucas D, Hidalgo A, et al. Bone marrow CD169+ macrophages promote the retention of hematopoietic stem and progenitor cells in the mesenchymal stem cell niche. J. Exp. Med. 2011;208(2):261–271.

41. Winkler IG, Sims NA, Pettit AR, et al. Bone marrow macrophages maintain hematopoietic stem cell (HSC) niches and their depletion mobilizes HSCs. Blood. 2010;116(23):4815–4828.

42. Zhou BO, Yu H, Yue R, et al. Bone marrow adipocytes promote the regeneration of stem cells and haematopoiesis by secreting SCF. Nat. Cell Biol. 2017;19(8):891–903.

43. Sykes SM, Scadden DT. Modeling human hematopoietic stem cell biology in the mouse. Semin. Hematol. 2013;50(2):92–100.

44. Leenaars CHC, Kouwenaar C, Stafleu FR, et al. Animal to human translation: a systematic scoping review of reported concordance rates. J. Transl. Med. 2019;17(1):223.

45. Watchman CJ, Bourke VA, Lyon JR, et al. Spatial distribution of blood vessels and CD34+ hematopoietic stem and progenitor cells within the marrow cavities of human cancellous bone. J. Nucl. Med. Off. Publ. Soc. Nucl. Med. 2007;48(4):645–654.

46. Bourke VA, Watchman CJ, Reith JD, et al. Spatial gradients of blood vessels and hematopoietic stem and progenitor cells within the marrow cavities of the human skeleton. Blood. 2009;114(19):4077–4080.

47. Takaku T, Malide D, Chen J, et al. Hematopoiesis in 3 dimensions: human and murine bone marrow architecture visualized by confocal microscopy. Blood. 2010;116(15):e41–55.

48. Flores-Figueroa E, Varma S, Montgomery K, Greenberg PL, Gratzinger D. Distinctive contact between CD34+ hematopoietic progenitors and CXCL12+ CD271+ mesenchymal stromal cells in benign and myelodysplastic bone marrow. Lab. Investig. J. Tech. Methods Pathol. 2012;92(9):1330–1341.

49. Guezguez B, Campbell CJV, Boyd AL, et al. Regional localization within the bone marrow influences the functional capacity of human HSCs. Cell Stem Cell. 2013;13(2):175–189.

50. Aguilar-Navarro AG, Meza-León B, Gratzinger D, et al. Human Aging Alters the Spatial Organization between CD34+ Hematopoietic Cells and Adipocytes in Bone Marrow. Stem Cell Rep. 2020;15(2):317–325.

51. Bauer M, Vaxevanis C, Al-Ali HK, et al. Altered Spatial Composition of the Immune Cell Repertoire in Association to CD34+ Blasts in Myelodysplastic Syndromes and Secondary Acute Myeloid Leukemia. Cancers. 2021;13(2):E186.

52. Kristensen HB, Andersen TL, Patriarca A, et al. Human hematopoietic microenvironments. PloS One. 2021;16(4):e0250081.

53. Tjin G, Flores-Figueroa E, Duarte D, et al. Imaging methods used to study mouse and human HSC niches: Current and emerging technologies. Bone. 2019;119:19–35.

54. Patel SS, Rodig SJ. Overview of Tissue Imaging Methods. Methods Mol. Biol. Clifton NJ. 2020;2055:455–465.

55. Patel SS, Lipschitz M, Pinkus GS, et al. Multiparametric in situ imaging of NPM1-mutated acute myeloid leukemia reveals prognostically-relevant features of the marrow microenvironment. Mod. Pathol. Off. J. U. S. Can. Acad. Pathol. Inc. 2020;33(7):1380–1388.

56. Walters DK, Jelinek DF. Multiplex Immunofluorescence of Bone Marrow Core Biopsies: Visualizing the Bone Marrow Immune Contexture. J. Histochem. Cytochem. Off. J. Histochem. Soc. 2020;68(2):99–112.

57. Coutu DL, Kokkaliaris KD, Kunz L, Schroeder T. Multicolor quantitative confocal imaging cytometry. Nat. Methods. 2018;15(1):39–46.

58. Berry S, Giraldo NA, Green BF, et al. Analysis of multispectral imaging with the AstroPath platform informs efficacy of PD-1 blockade. Science. 2021;372(6547):eaba2609.

59. Naveiras O, Nardi V, Wenzel PL, et al. Bone-marrow adipocytes as negative regulators of the haematopoietic microenvironment. Nature. 2009;460(7252):259–263.

60. Pang WW, Price EA, Sahoo D, et al. Human bone marrow hematopoietic stem cells are increased in frequency and myeloid-biased with age. Proc. Natl. Acad. Sci. U. S. A. 2011;108(50):20012–20017.

61. Lengefeld J, Cheng C-W, Maretich P, et al. Cell size is a determinant of stem cell potential during aging. Sci. Adv. 2021;7(46):eabk0271.

62. Heazlewood SY, Ahmad T, Cao B, et al. High ploidy large cytoplasmic megakaryocytes are hematopoietic stem cells regulators and essential for platelet production. Nat. Commun. 2023;14(1):2099.

63. Poscablo DM, Worthington AK, Smith-Berdan S, Forsberg EC. Megakaryocyte progenitor cell function is enhanced upon aging despite the functional decline of aged hematopoietic stem cells. Stem Cell Rep. 2021;16(6):1598–1613.

64. Sanjuan-Pla A, Macaulay IC, Jensen CT, et al. Platelet-biased stem cells reside at the apex of the haematopoietic stem-cell hierarchy. Nature. 2013;502(7470):232–236.

65. Grover A, Sanjuan-Pla A, Thongjuea S, et al. Single-cell RNA sequencing reveals molecular and functional platelet bias of aged haematopoietic stem cells. Nat. Commun. 2016;7:11075.

66. Radtke AJ, Postovalova E, Varlamova A, et al. A Multi-scale, Multiomic Atlas of Human Normal and Follicular Lymphoma Lymph Nodes. 2022;2022.06.03.494716.

67. Zhao J, Ghimire A, Liesveld J. Marrow failure and aging: The role of “Inflammaging.” Best Pract. Res. Clin. Haematol. 2021;34(2):101283.

68. Patel SS, Weirather JL, Lipschitz M, et al. The microenvironmental niche in classic Hodgkin lymphoma is enriched for CTLA-4-positive T cells that are PD-1-negative. Blood. 2019;134(23):2059–2069.

69. Pan S, Hu R, Long G, et al. Adversarially Regularized Graph Autoencoder for Graph Embedding. 2019;

