## Supplemental Data for "Single Cell Resolution Spatial Mapping of Human Hematopoiesis Reveals Aging-Associated Topographic Remodeling"

### Supplemental Table 1

| Case ID | Age (years) | M/F | BMI (kg/m2) | Hb (g/dL) | MCV (fL) | ANC (cells/microliter) | Plts (K cells/microliter) | MPV (fL) | M:E (Aspirate diff) | Blast cells (Aspirate diff) | Promyelocytes (asp) | Proerythroblast (asp) | plasma cells (asp) | Lymphs (MFC) | monos/grans (MFC) | plasma cells (MFC) | Karyotype |
|---|---|---|---|---|---|---|---|---|---|---|---|---|---|---|---|---|---|
| WCM62 | 5 | M | 14 | 11.6 | 76 | 5002 | 373 | 7.3 | 3.6 | 3% | 5.00% | 2.00% | 1.00% | 7.30% | 76.10% | N/A | 46,XY[20] |
| WCM59 | 10 | F | 15.7 | 10.9 | 70.3 | 13700 | 461 | 11.1 | N/A | N/A | N/A | N/A | N/A | N/A | N/A | N/A | N/A |
| WCM60 | 10 | M | 16.5 | 12.9 | 81.9 | 10656 | 403 | 8.9 | 4.7 | 3% | 7.00% | 1.00% | 3.00% | 12.50% | 81.50% | N/A | 46,XY[20] |
| WCM66 | 15 | F | 26.93 | 7.8 | 72.8 | 1836 | 71 | 9.8 | 2.7 | 1% | 2.00% | 1.00% | 1.00% | 10.80% | 86.50% | 0.169% | 46,XX[20] |
| WCM68 | 15 | F | 18.24 | 12.5 | 81.4 | 8500 | 359 | 8.5 | N/A | N/A | N/A | N/A | N/A | N/A | N/A | N/A | N/A |
| WCM70 | 15 | F | N/A | 9.5 | 72.9 | 11684 | 341 | 8.7 | N/A | 2% | 1.00% | 1.00% | 0.00% | 2.90% | 92.80% | N/A | 46,XX[20] |
| WCM69 | 16 | F | 20.4 | 10.5 | 78.7 | 19600 | 400 | 8.4 | 6.1 | 2% | 4.00% | 0.00% | 2.00% | N/A | N/A | N/A | N/A |
| WCM65 | 18 | M | 29.26 | 12 | 82.5 | 3970 | 315 | 9.2 | 2.7 | 2% | 4.00% | 2.00% | 1.00% | 7.20% | 90.50% | 0.100% | 46,XY[20] |
| WCM67 | 18 | F | 23.5 | 9.9 | 75.8 | 4237 | 378 | 8.8 | 3.9 | 3% | 6.00% | 2.00% | 2.00% | 5.10% | 92.70% | 0.046% | 46,XX[20] |
| WCM61 | 19 | M | 20.43 | 14.4 | 84 | 7900 | 284 | 8.4 | N/A | N/A | N/A | N/A | N/A | N/A | N/A | N/A | N/A |
| WCM64 | 19 | M | 22.81 | 15.6 | 87.1 | 5100 | 188 | 8.4 | 1.6 | 3% | 1.00% | 1.00% | 0.00% | 3.80% | N/A | N/A | 46,XY[20] |
| WCM10 | 28 | M | 26.04 | 12.4 | 79.7 | 5915 | 501 | 8.1 | 2.3 | 1% | 0.00% | 1.00% | 2.00% | 8.50% | 88.10% | N/A | 46,XY[20] |
| WCM9 | 37 | F | 22.84 | 11.6 | 85.8 | 2860 | 251 | 7 | 2.3 | 3% | 1.00% | 1.00% | 3.00% | 10.40% | 81.80% | 0.058% | 46,XX,inv(9)(p12q13)c[20] |
| WCM38 | 41 | M | 23.99 | 13.2 | 87.1 | 2400 | 147 | 10.2 | 3.3 | 2% | 1.00% | 1.00% | 1.00% | 5.50% | 87.90% | N/A | 46,XY[20] |
| WCM11 | 50 | M | 34.18 | 15.7 | 87.9 | 6912 | 283 | 7.5 | 3 | 4% | 6.00% | 2.00% | 6.00% | 16.00% | 81.10% | N/A | 46,XY[20] |
| WCM8 | 52 | F | 30.98 | 11.9 | 84.5 | 3960 | 257 | 7.4 | 2 | 3% | 4.00% | 2.00% | 6.00% | 23.70% | 73.10% | N/A | 46,XX[20] |
| WCM35 | 61 | F | 27.48 | 11.1 | 108 | 2478 | 242 | 6 | 1.5 | 2% | 0.00% | 2.00% | 3.00% | 14.10% | 80.90% | N/A | 46,XX[20] |
| WCM31 | 70 | F | 35.12 | 13.2 | 93.5 | 3600 | 164 | 9.1 | 2.4 | 2% | 1.00% | 1.00% | 1.00% | 3.50% | 90.80% | N/A | 46,XX[20] |
| WCM30 | 71 | M | 25.6 | 16.5 | 94.9 | 4292 | 273 | 8.4 | 2.7 | 3% | 5.00% | 2.00% | 6.00% | 10.90% | 84.20% | N/A | 46,XY[20] |
| WCM34 | 74 | F | 24.92 | 12.6 | 83.6 | 5112 | 180 | 8.5 | 2.2 | 1% | 4.00% | 2.00% | 4.00% | 12.90% | 84.10% | 0.005% | 46,XX[20] |
| WCM1 | 75 | M | 30.28 | 14.1 | 97 | 4216 | 153 | 7.9 | 2.3 | 3% | 2.00% | 2.00% | 0.00% | N/A | N/A | N/A | N/A |
| WCM33 | 76 | M | 24.89 | 11.7 | 98.1 | 5200 | 383 | 8.7 | 7 | 3% | 6.00% | 1.00% | 14.00% | 8.00% | 88.10% | 0.500% | 46,XY[20] |
| WCM55 | 80 | F | 26.45 | 8 | 87.1 | 2597 | 253 | 7.6 | 1.3 | 1% | 2.00% | 2.00% | 5.00% | 12.10% | 84.90% | N/A | 46,XX[20] |
| WCM56 | 80 | M | 27.4 | 12.4 | 82.9 | 4560 | 241 | 7.7 | 2.7 | 1% | 2.00% | 1.00% | 1.00% | 16.20% | 80.40% | 0.020% | 46,XY[20] |
| WCM36 | 83 | M | 28.62 | 10.2 | 94.7 | 5313 | 205 | 7.8 | 2.5 | 0% | 2.00% | 4.00% | 5.00% | 9.90% | 88.00% | 0.100% | 46,XY[20] |
| WCM53 | 84 | M | 16.51 | 9.6 | 78.8 | 1827 | 204 | 7.7 | 2.9 | 3% | 6.00% | 2.00% | 7.00% | 6.30% | 84.00% | N/A | 46,XX[20] |
| WCM32 | 86 | M | 23.7 | 12.3 | 82.1 | 2142 | 193 | 8.2 | 2.2 | 2% | 3.00% | 2.00% | 17.00% | 29.40% | 66.40% | 0.600% | 45,X,-Y[3]/46,XY[17] |
| WCM54 | 87 | F | 21.06 | 11.1 | 89.7 | 12056 | 170 | 9.5 | 2.4 | 2% | 1.00% | 1.00% | 1.00% | 8.60% | 86.00% | 0.200% | 46,XX[20] |
| WCM57 | 89 | M | 28.81 | 13 | 95.3 | 10004 | 119 | 12.2 | 2.4 | 1% | 2.00% | 2.00% | 5.00% | 6.70% | 92.40% | 0.030% | 45,X,-Y[8]/46,XY[12] |

M, Male; F, Female; BMI, Body Mass Index; Hb, Hemoglobin (g/dL); MCV, Mean Corpuscular Volume (fL), ANC, Absolute Neutrophil Count; Plts, Platelet Count (thousand/microliter); MPV, Mean Platelet Volume (fL)

Asp, Aspirate Smear Differential Count; MFC, Multiparameter Flow Cytometry; M:E, Myeloid to Erythroid Ratio; Monos/Grans, Monocytes + Granulocytes

### Supplemental Figure 1

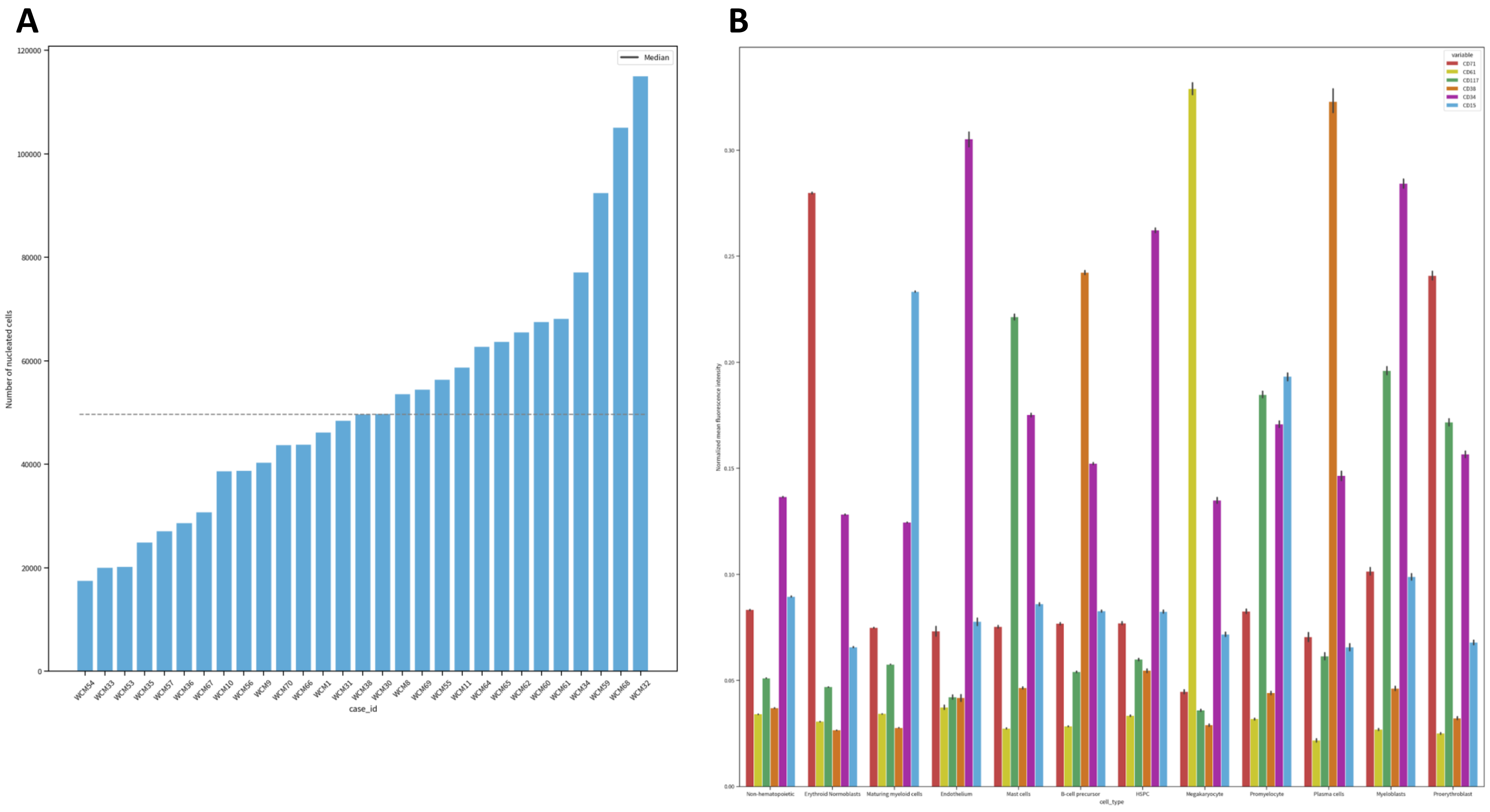


**Supplemental Figure 1. Total nucleated cell counts and MFI marker values for analyzed antigens for each cell type. A.** A total of 1,510,295 nucleated cells were analyzed across the entire cohort (median: 49,669 cells/sample, range: 17,543 - 115,027). **B.** Bar plot with normal mean fluorescence intensity (MFI) values (y-axis) for each antigen tested associated with each defined cell type (x-axis) including non-hematopoietic elements (NHEs), erythroid normoblasts, maturing myeloid cells (MMCs), endothelial cells (endothelium), mast cells, likely B-cell precursors, HSPCs, megakaryocytes, promyelocytes, plasma cells, myeloblasts, and proerythroblasts.

### Supplemental Figure 2

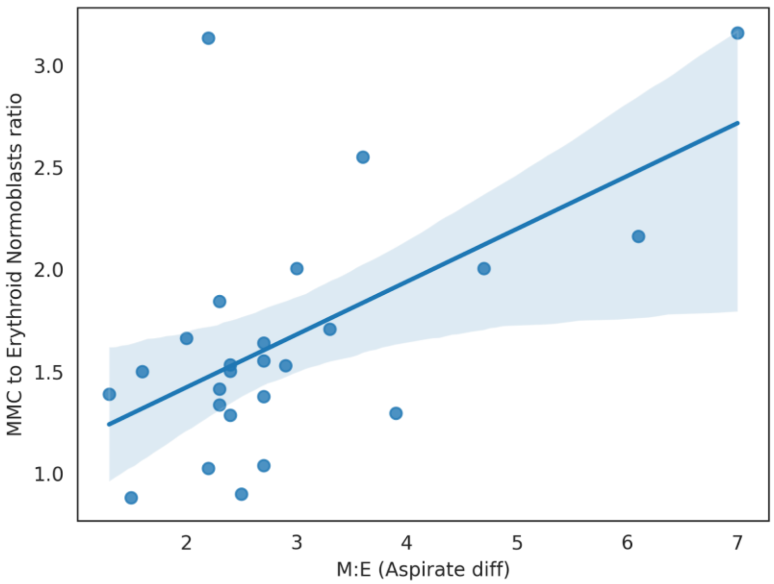


**Supplemental Figure 2.** Positive correlation identified for comparison of ratio of MMCs to erythroid normoblasts (y-axis) with myeloid to erythroid cell count ratio as determined by manual aspirate smear differential counting (x-axis) [Pearson r = 0.58, spearman r = 0.45, p=0.025].

### Supplemental Figure 3

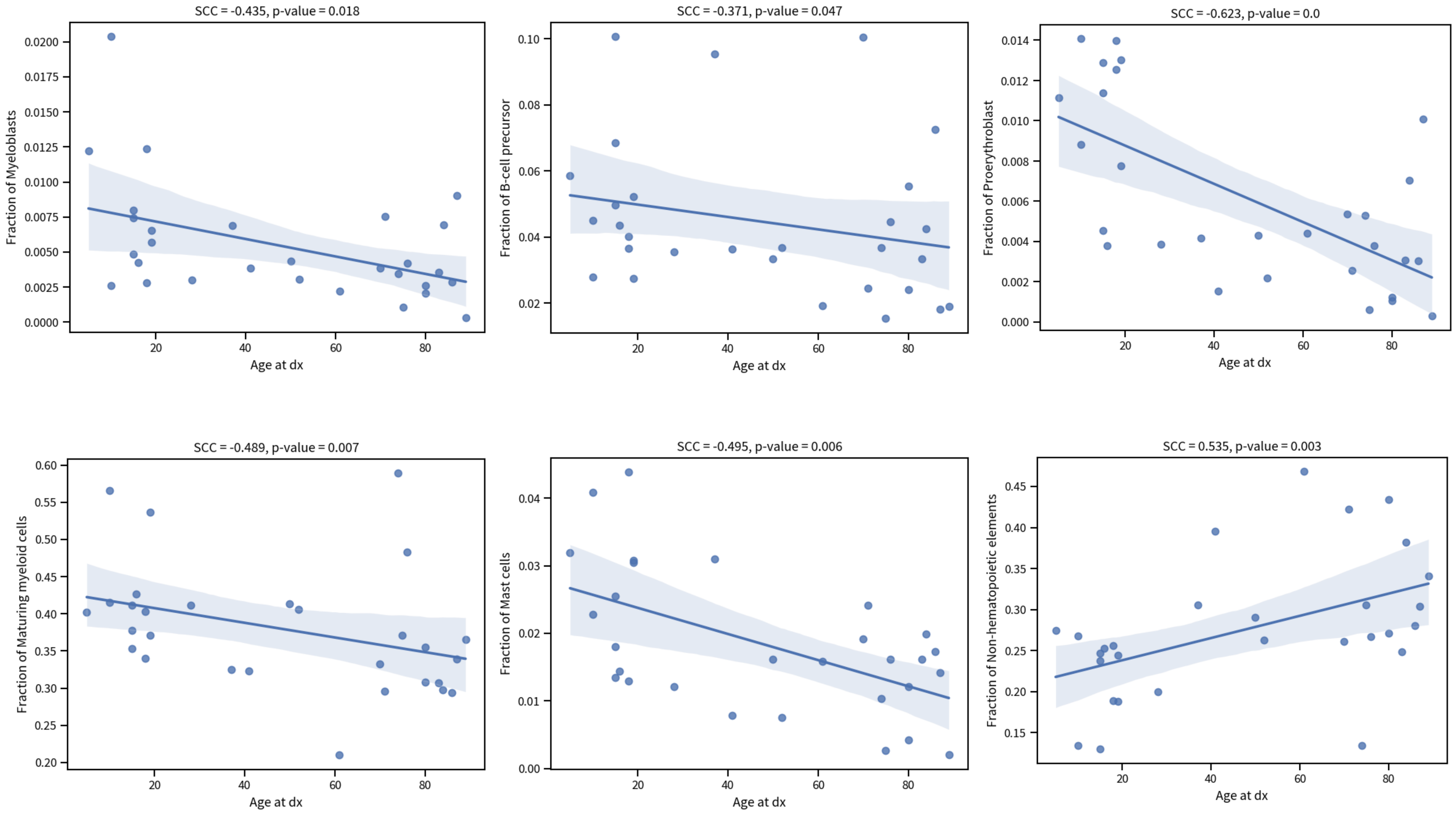


**Supplemental Figure 3. Correlations of cell type proportions with increasing patient age**: myeloblasts (spearman r = -0.435, p = 0.018), likely B-cell precursors (spearman r = -0.371, p = 0.047), proerythroblasts (spearman r = -0.623, p < 0.01), MMCs (spearman r = -0.489, p = 0.007), mast cells (spearman r = -0.495, p = 0.006), and non-hematopoietic elements (spearman r = -0.535, p = 0.003).

### Supplemental Figure 4

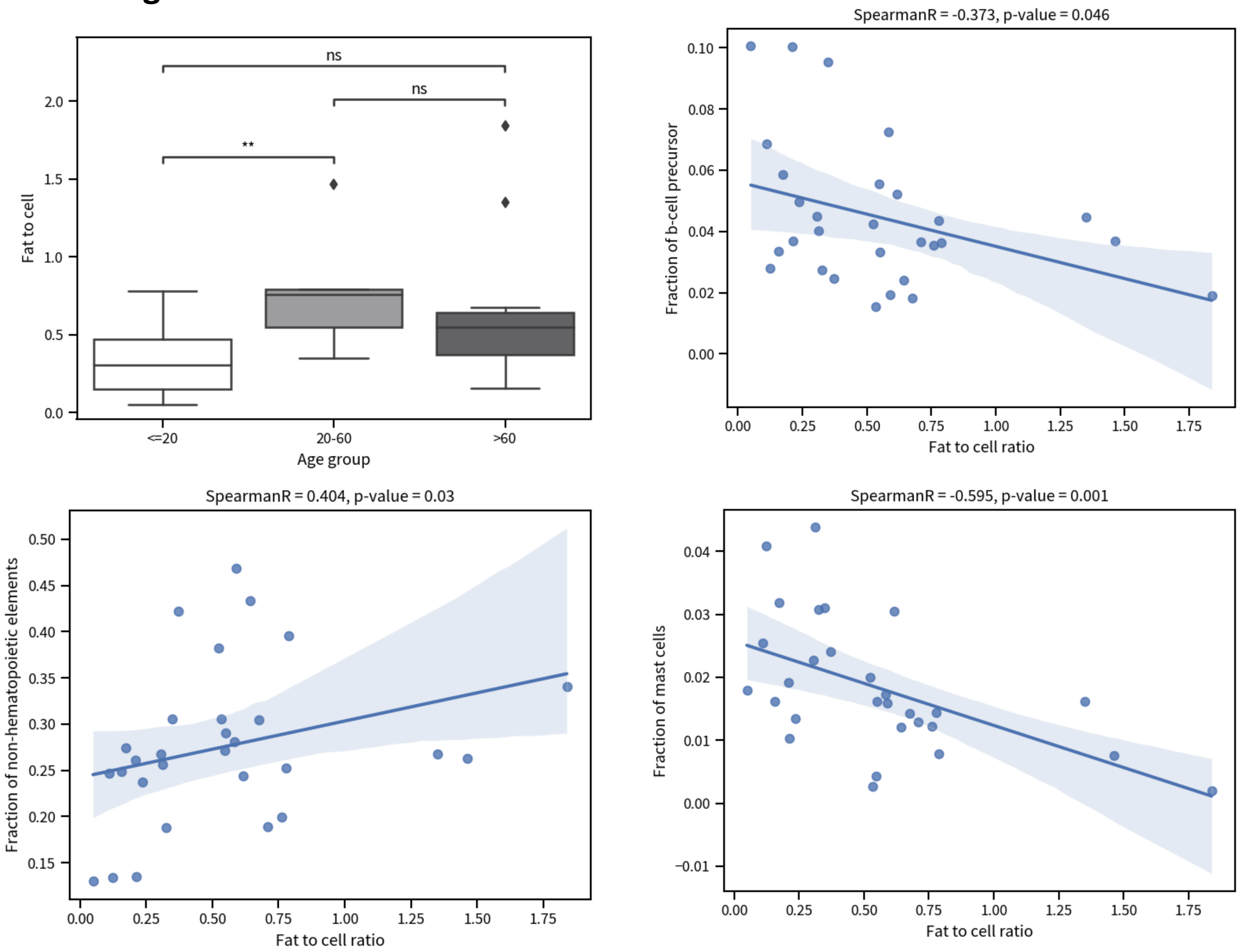


**Supplemental Figure 4. Additional associations with increasing fat:cell area ratio.** Significant difference in fat:cell area ratio between <20 years and 20-60 years marrows (0.34±0.25 vs. 0.78±0.42, p = 0.009; p=0.058 for <=20 vs >60), but no difference between extremes of age. Increasing fat:cell area ratio correlates with proportions of likely B-cell precursors (spearman r = -0.373, p = 0.046), NHEs (spearman r = 0.404, p = 0.03), and mast cells (spearman r = -0.595, p = 0.001).

### Supplemental Figure 5

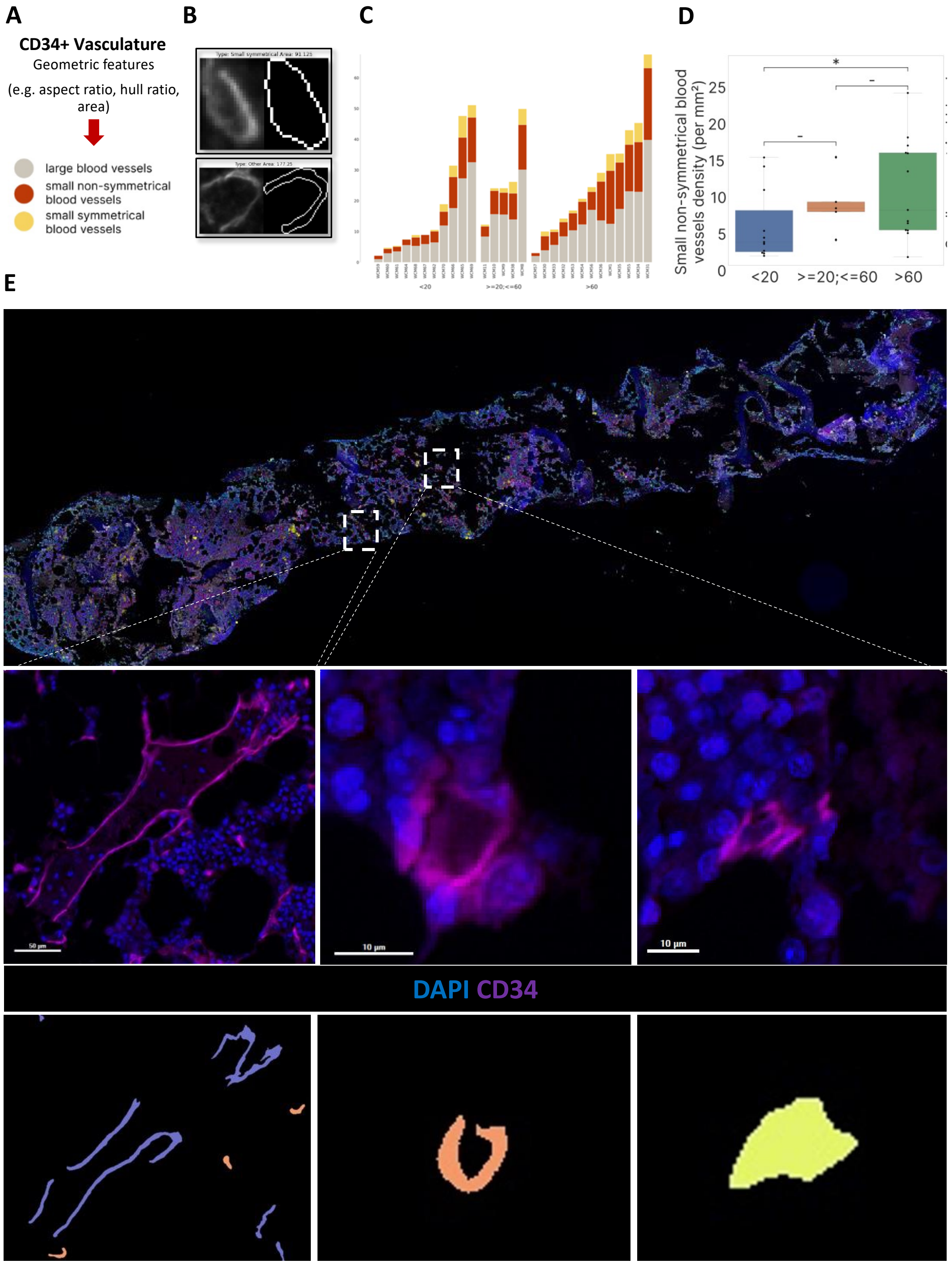


**Supplemental Figure 5. Assessment of overall vascular density and CD34+ vascular morphologies. A.** Separation of CD34+ endothelial-lined vascular elements into three morphologic subtypes (large, small symmetrical, and small non-symmetrical) based on geometric features including aspect ratio, hull ratio, and area. **B.** Representative vascular masks for small symmetrical (top) and small non-symmetrical (bottom) vessel subtypes. **C.** Heterogeneous distribution of vascular density morphology within age groups, with no significant difference in overall vascular density observed between age groups. **D.** Density of small non-symmetrical vessels is greater in >60 years individuals vs. ≤20 years individuals (0.7±0.68 mm vs. 1.1±0.95 mm, p=0.013). **E.** Representative MxIF and digitized vascular mask images of large, small symmetrical, and small non-symmetrical vessel types (left to right).

### Supplemental Figure 6

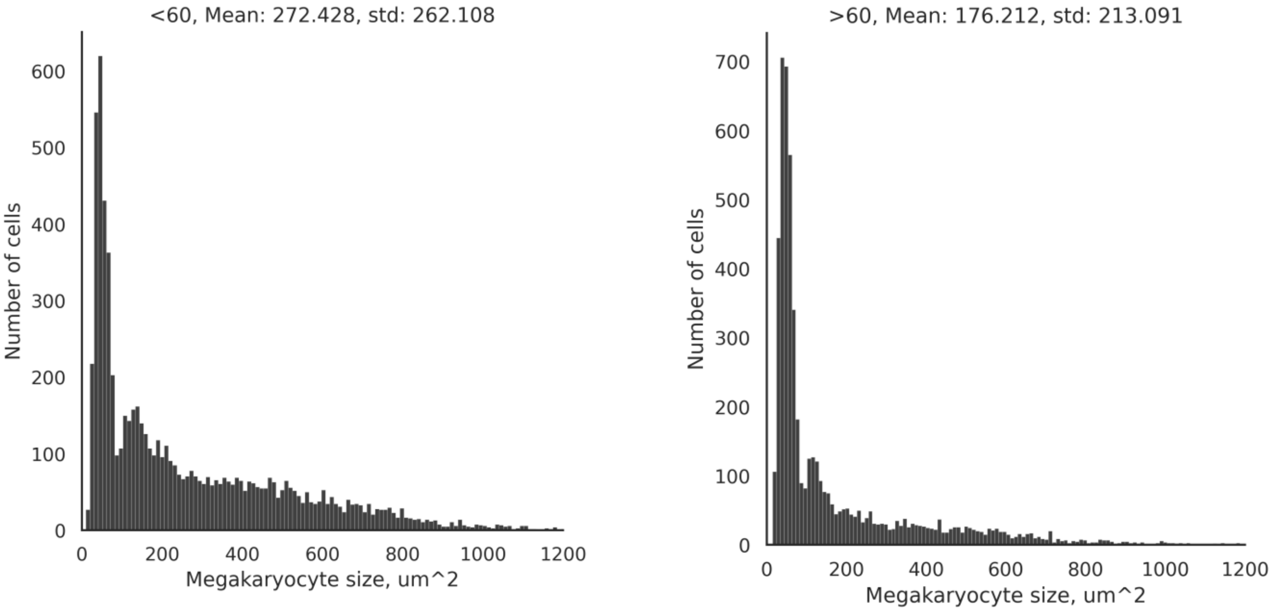


**Supplemental Figure 6. Differences in megakaryocyte size distribution between youngest and oldest bone marrows.**

### Supplemental Figure 7

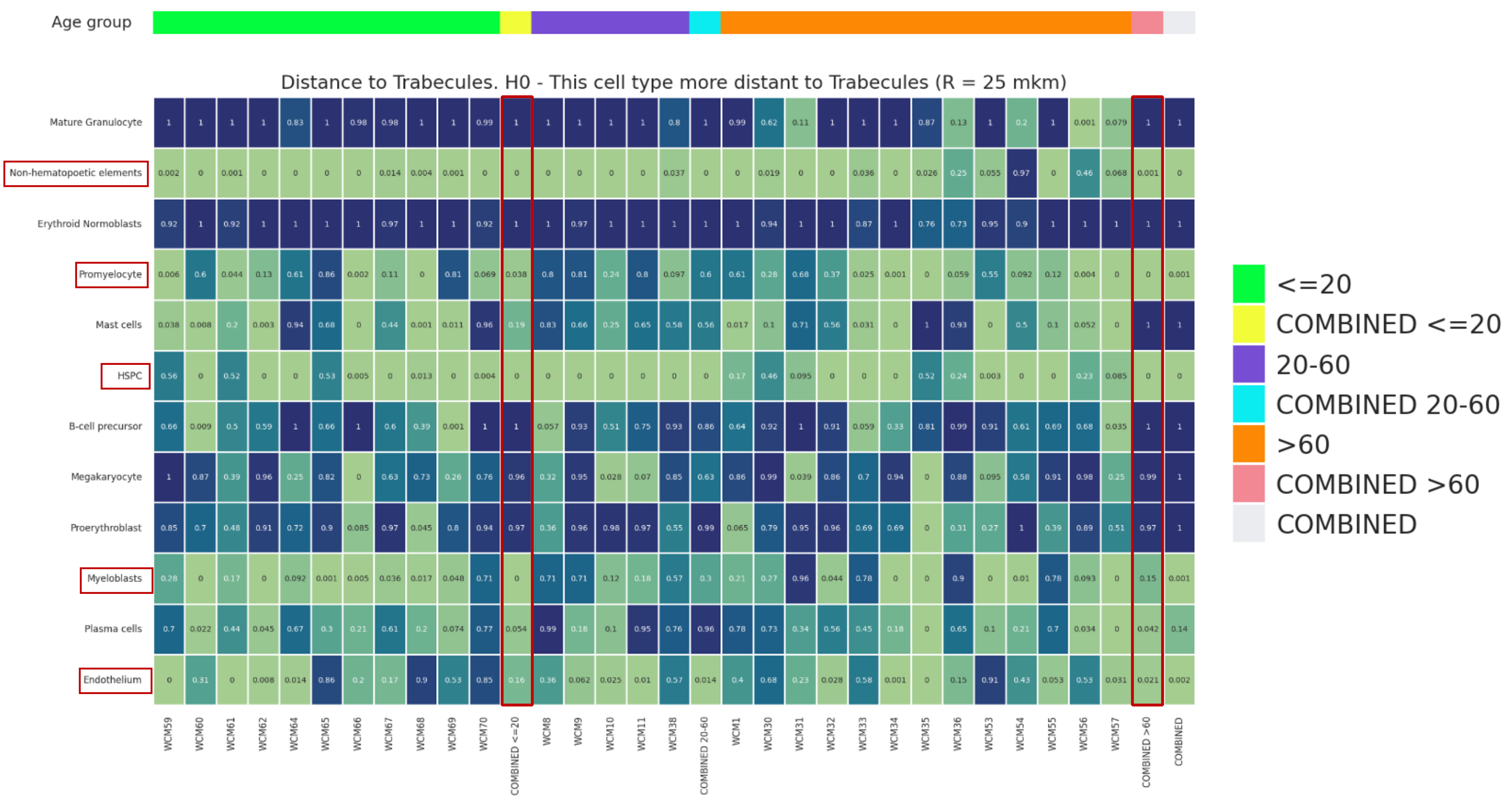


**Supplemental Figure 7. Permutation testing results for comparison of observed versus randomized distribution for all cell types, with assessment of cell type enrichment within 25 microns of bone trabeculae.** Each column indicates an individual specimen, while columns marked yellow (≤20 years) and peach (>60 years) denote analyses of entire age groups. Specimens grouped by increasing age. Boxes color-coded to reflect trend toward statistically significant enrichment (lightest green) or lack of proximity relationship (dark blue). Numbers within boxes are p-values. Analysis revealed significant proximity association to bone trabeculae for HSPCs, promyelocytes, and non-hematopoietic elements in ≤20 years and >60 years age groups. Myeloblasts are also proximal to bone in ≤20 years marrows, while >60 years marrows show relative enrichment for plasma cells and endothelium. Other cell types, including proerythroblasts, megakaryocytes, MMCs, and erythroid normoblasts are consistently distant from bone trabeculae in both age groups.

### Supplemental Figure 8

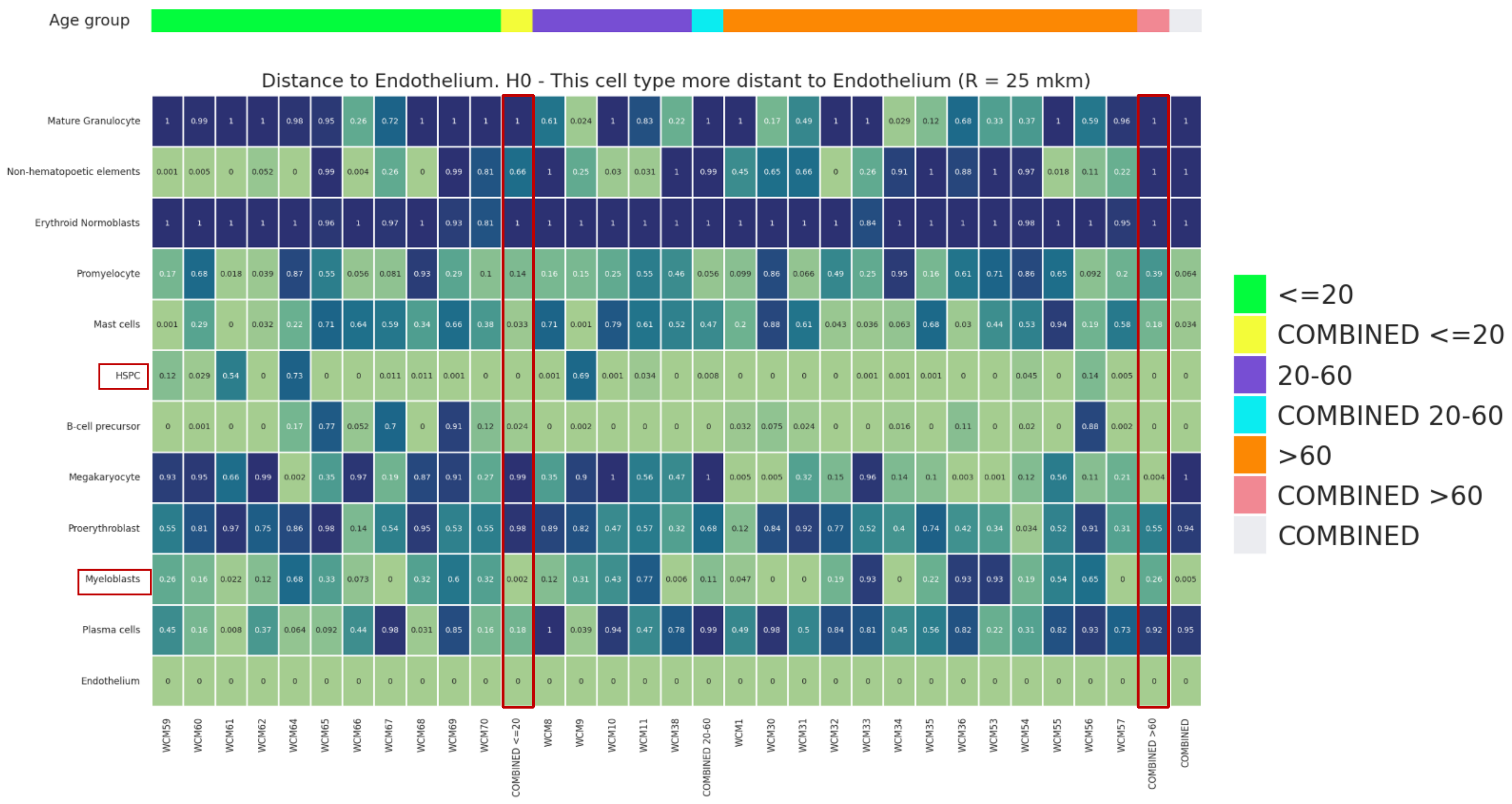


**Supplemental Figure 8. Permutation testing results for comparison of observed versus randomized distribution for all cell types, with assessment of cell type enrichment within 25 microns of CD34+ vasculature.** Each column indicates an individual specimen, while columns marked yellow (≤20 years) and peach (>60 years) denote analyses of entire age groups. Specimens grouped by increasing age. Boxes color-coded to reflect trend toward statistically significant enrichment (lightest green) or lack of proximity relationship (dark blue). Numbers within boxes are p-values. Analysis revealed significant proximity association to CD34+ vasculature for HSPCs and likely B-cell precursors (hematogones) in ≤20 years and >60 years age groups. Myeloblasts and mast cells are also proximal to CD34+ vasculature in ≤20 years marrows, while >60 years marrows show relative enrichment for megakaryocytes. Other cell types, including erythroid normoblasts and MMCs are consistently distant from CD34+ vasculature in both age groups. ≤20 marrows exhibit a more pronounced distancing of proerythroblasts, while >60 years exhibit a similar distancing for plasma cells.

### Supplemental Figure 9

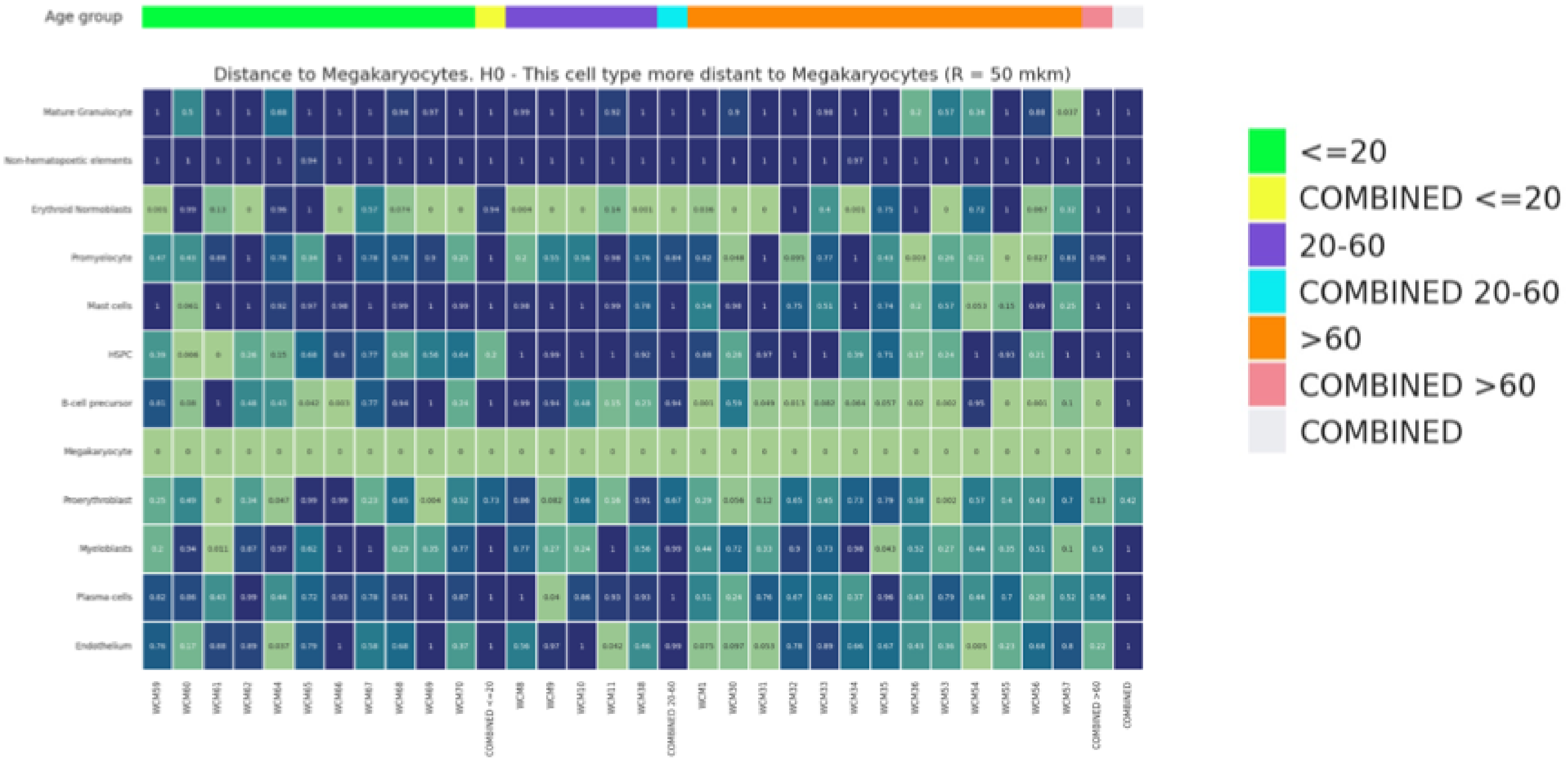


**Supplemental Figure 9. Permutation testing results for comparison of observed versus randomized distribution for all cell types, with assessment of cell type enrichment within 50 microns of megakaryocytes.** Each column indicates an individual specimen, while columns marked yellow (≤20 years) and peach (>60 years) denote analyses of entire age groups. Specimens grouped by increasing age. Boxes color-coded to reflect trend toward statistically significant enrichment (lightest green) or lack of proximity relationship (dark blue). Numbers within boxes are p-values. Analysis revealed relative proximity association between HSPCs and megakaryocytes in ≤20 years ($p=0.20$) versus >60 years ($p=1.0$) age groups. Likely B-cell precursors were enriched near megakaryocytes in >60 years marrows ($p<0.01$) but not in young and middle age groups. No other significant associations were identified. All other cell types were found to be non-enriched near megakaryocytes in all age groups.

### Supplemental Figure 10

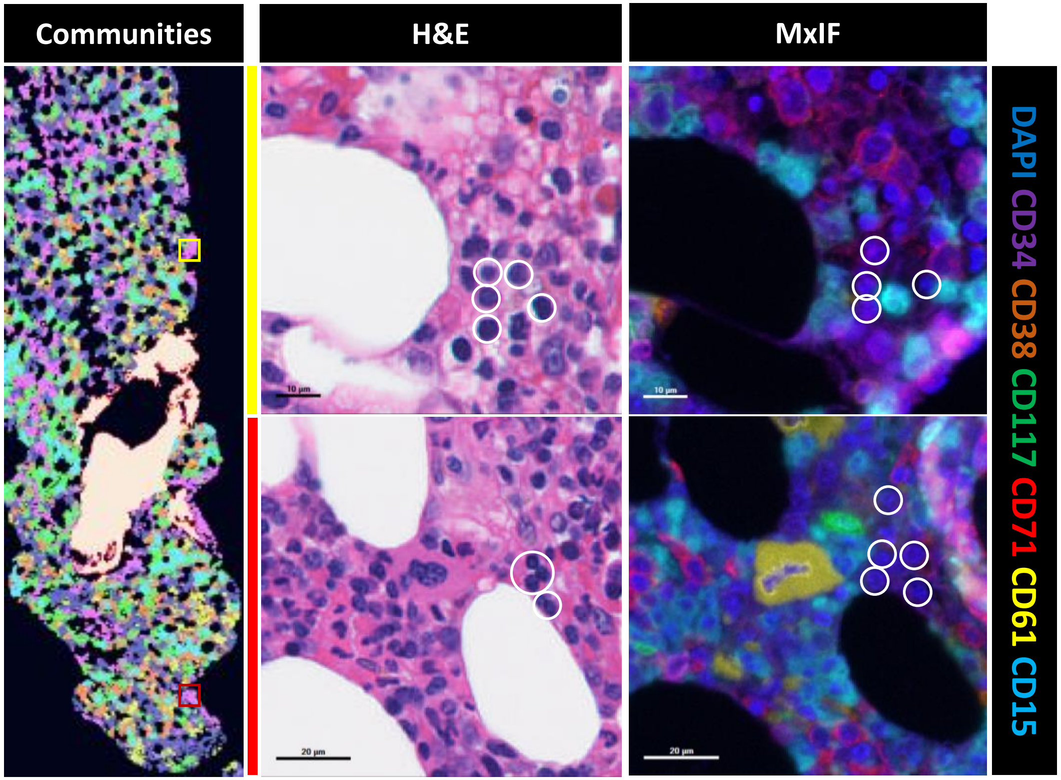


**Supplemental Figure 10. Detection of Cluster 1 foci in paired H&E and MxIF images.** From left to right: community/cluster map (first column), H&E images from same case (middle column), MxIF images from same case (third column). Representative Cluster 1 foci in the community map are highlighted with yellow and red boxes. Yellow region further investigated in top row of H&E/MxIF images, red region in bottom row. Representative cells with cytomorphologic features of mature lymphocytes on H&E staining have phenotype on MxIF corresponding to NHEs (DAPI+/negative for all tested antibodies).
